# The diet-microbiome axis instructs gut barrier integrity by modulating colonocyte JNK-P38 duality

**DOI:** 10.1101/2025.01.24.634634

**Authors:** Chen Chongtham, Trisha Biswas, Namaste Kumari, Raunak Kar, Jyotsna, S Jayalakshmi, Archana Pant, Veena S. Patil, Gopalakrishnan Aneeshkumar Arimbasseri

**Affiliations:** Molecular Genetics Laboratory, National Institute of Immunology, New Delhi, India; Immunogenomics Laboratory, National Institute of Immunology, New Delhi, India; Immunometabolism Laboratory, National Institute of Immunology, New Delhi, India

**Author notes:** **Corresponding Author:** Gopalakrishnan Aneeshkumar Arimbasseri, Molecular Genetics Laboratory, National Institute of Immunology, Aruna Asaf Ali Marg, New Delhi 110067. These authors contributed equally (Trisha Biswas, Namaste Kumari). The authors have declared that no conflict of interest exists.

**Keywords:** Weaning, JNK2 pathway, P38 pathway, Gut microbiome, Gut barrier, Inflammation

## Abstract

Weaning involves a nutritional shift from fat-rich milk to carbohydrate-based solid food, reshaping metabolism, microbiota, and gut immune tolerance. While dairy remains a component of the human diet beyond weaning, the impact of continued milk supplementation on gut epithelial homeostasis remains poorly understood. Here, using a mouse model, we show that continued milk-based feeding post-weaning promotes intestinal barrier function by enriching the commensal bacterium *Dubosiella newyorkensis*, which produces acetate to activate epithelial JNK2 signaling. This pathway enhances barrier integrity and suppresses inflammation induced by mild Dextran Sodium Sulphate (DSS) treatment. In contrast, feeding a lard-based high-fat diet or transient pharmacologic inhibition of JNK2 induces epithelial P38 activation, resulting in barrier disruption and inflammation. Importantly, the beneficial effects of milk were observed only if initiated during the weaning period, when the microbiome is in a metastable transitional state. Initiation of the same intervention two weeks after weaning led to P38 activation and inflammatory responses. These findings highlight the microbiome-dependent, stage-specific effects of diet and underscore the importance of early-life nutritional interventions for long-term gut health.

## Introduction

The diet-microbiome-host axis in the gut has emerged as a key factor in chronic diseases, including inflammatory bowel diseases and metabolic and cardiovascular diseases. While several diets, such as the Mediterranean diet, fermented food, and high-fibre diet, have been considered microbiome-friendly, others, including high-fat and western diets, promote gut dysbiosis, causing gut inflammation (*1*). Microbiome-directed dietary interventions have been an effective tool to fight childhood malnutrition (*2*). However, to formalise the nature of different diet-microbiome-gut interactions and develop novel intervention strategies, it is essential to identify the molecular signatures and mechanisms that drive such interactions.

During the postnatal period, several milk components, including Immunoglobulin A, complex oligosaccharides, higher fat content, etc., are essential for developing a healthy microbiome (*3*). The weaning period is one of the most critical stages of microbiome maturation in mammals, with the changes in microbiota during this period having long-lasting effects on gut and metabolic health (*4–6*). During the weaning period, the changes in the diet lead to the establishment of the adult microbiome, which is associated with a spike in inflammation, followed by its resolution (*4*). This ‘weaning reaction’ is essential for developing immune tolerance towards the altered microbiome. Interrupting this process leads to increased susceptibility to DSS-induced gut inflammation.

Humans consume milk and milk products even after weaning, unlike most other animals. Indeed, dairy products are a significant source of healthy nutrition for humans. However, very little is known about the effect of continued milk supplementation post-weaning on gut function. While patients with inflammatory bowel disease tend to exclude milk and milk products, conflicting conclusions have been made on the effect of milk and dairy products on gut inflammation (*7–9*). The microbiome plays an important role in diet-induced gut pathologies. Recent evidence indicates that diet is the primary determinant of the microbial ecosystem within the gut (*10*). However, it is possible that the trajectory of a given diet in shaping the microbiome and inflammatory landscape of the gut could be dependent on the already established microbiome. Such context-dependent differences in the pre-existing gut-microbiome composition and function could be one of the driving forces determining milk’s positive and negative effects on gut health.

Host-microbiome interactions are controlled by a multitude of factors, including microbial metabolites such as short-chain fatty acids, host-secreted proteins such as antimicrobial peptides, metabolites, dietary components, etc. (*11*). The mechanisms by which these metabolites enhance gut barrier function or lower pathological inflammation are not entirely understood. JNK and P38 pathways are among the core signalling pathways in mammalian physiology, regulating various processes such as cell proliferation and inflammation. Both paths in the gut are known to affect inflammation, gut barrier function, epithelial regeneration, and even cancer (*12–14*). AP-1 transcription factors are one of the significant classes of targets for the JNK pathway, which are known to regulate gut regeneration and cancer. The JNK pathway was shown to enhance epithelial regeneration by enhancing WNT signalling (*15*). Deletion of MBD3, an inhibitory protein associated with unphosphorylated c-Jun, leads to enhanced progenitor proliferation (*16*, *17*). Genetic depletion experiments have shown that JNK2, but not JNK1, enhances gut barrier function and protection against dextran sodium sulfate treatment (DSS) (*18*). However, another study has demonstrated that osmotic stress-induced activation of JNK2 deteriorates tight junction protein expression and gut barrier function (*19*), indicating the context-dependent effects of JNK2 activation in the gut epithelium. On the other hand, ulcerative colitis and Crohn’s disease patient IECs have an elevated level of P38 (*20*), and inhibition of P38 reduces IBD (*21*). Activated P38 works downstream of the cytokine signalling pathways, such as TNF-α, IL-1β, and IL-8, which further enhance inflammation (*22*). However, how diet and gut microbiome alter these two crucial pathways in different contexts of gut inflammation is unknown.

Here, we show that post-weaning milk-based diet consumption, depending on the timing of the initiation, elicits opposite effects on the gut barrier integrity and susceptibility to inflammation in mice. Mice weaned directly onto milk-based diets maintain gut epithelial integrity even after a challenge with 1.5% DSS for 7 days, while a 2-week delay in initiation of the same diet elicits opposite effects. These opposing effects of the same diet in two contexts could be explained by the distinct trajectories of microbiome modulation depending on the timing of initiation of the diet. Molecular analysis reveals a complex liaison between the gut microbiome and the epithelial cells, which protects the epithelial layer from DSS insult, with the dichotomy between epithelial JNK2 and P38 pathways being the deciding factor in pathological inflammation. We identified that *D. newyorkensis* produces acetate, which mediates activation of JNK2 to protect gut barrier function.

## Results

### Post-weaning milk-based diets protect from mild DSS-induced gut inflammation

To address the effect of an extended milk-like diet on gut health, we weaned 24 days old pups onto one of two milk-based diets: a custom-designed, milk-fat-based diet (MFD) that mimics natural milk composition or a commercial milk-based formula, Lactogen 2 (MBD), that contains 54.8% milk solids. These two diets differ in macronutrient composition, with MFD providing higher fat content and Lactogen 2 supplying more carbohydrates. (Table S1, Figure S1A). These diets have previously been shown to rescue the growth and metabolic defects in whole-body vitamin D receptor knock-out mice (*23*). Initiating the milk-fat diet at 3 weeks of age did not affect the onset of weaning reaction (Figure S1B) (*4*). To distinguish the effect of milk components from the effect of high-fat content alone, we also included a lard-based high-fat diet (HFD), which is known to induce gut inflammation (Table S1, Figure S1A) (*24*). We fed mice these diets or a regular chow diet (RCD) for 4 weeks, and subsequently treated them with 1.5% DSS for 7 days to induce gut inflammation (Figure 1A).

**Figure 1:**
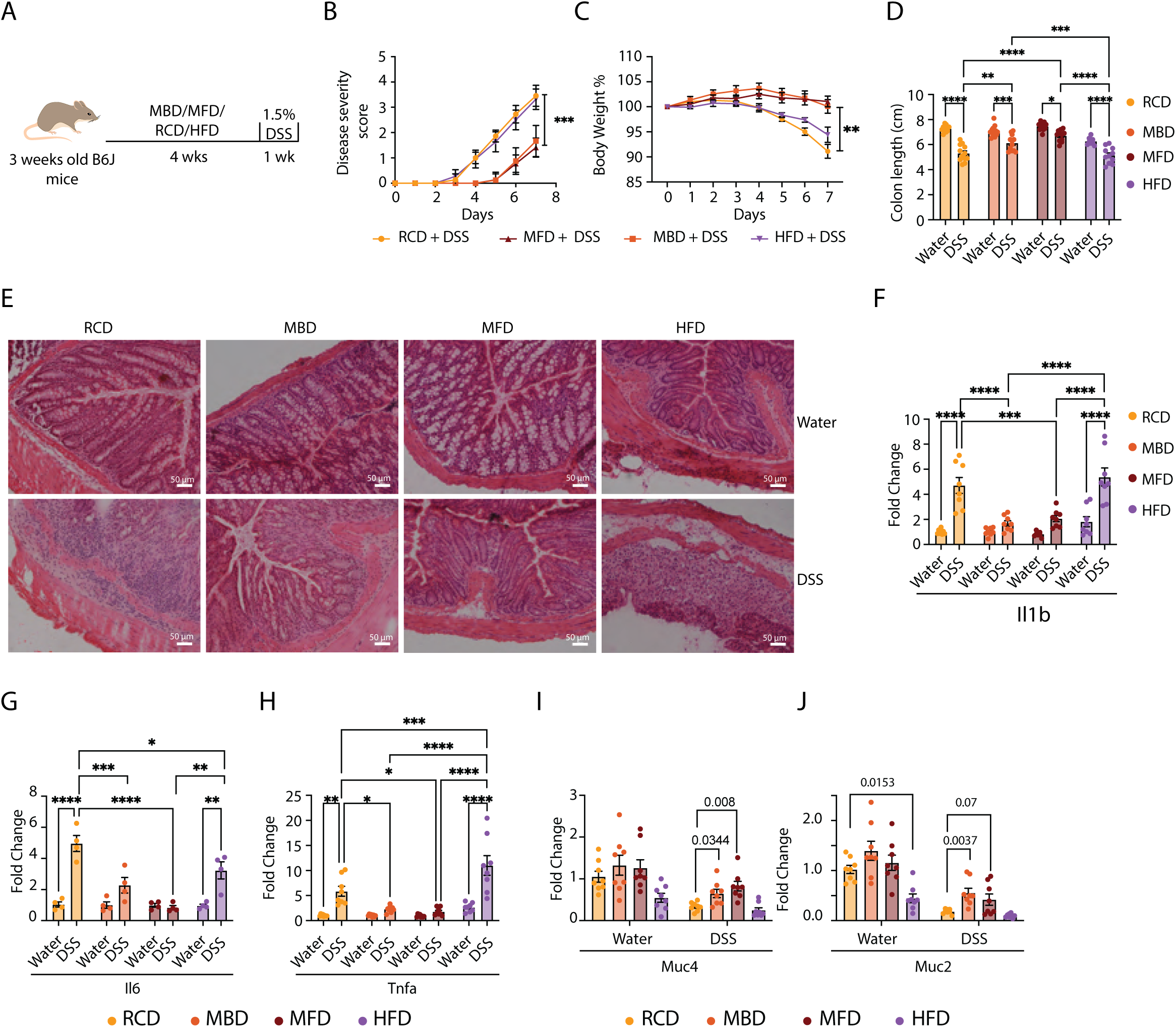
Post-weaning milk-based diets protect from DSS-induced gut inflammation. **A:** An Illustration of the DSS-induced mouse colitis experimental model. 3 weeks old mice were weaned to the indicated diets. After 4 weeks, they were subjected to 1.5% DSS treatment. **B:** Disease severity score calculated from the rectal bleeding and stool consistency for different experimental groups indicated (n = 12). **C:** Percentage of body weight after DSS treatment with day 0 as the baseline. Different experimental groups are indicated (n = 12). **D:** Colon length measured in centimeters for different experimental groups (n = 12). The images of the colon are given in Figure S1C. **E:** Representative images of H&E-stained colon tissue sections (DSS- or water-treated mice fed RCD, MBD, MFD, and HFD. (Scale bars: 50 μm). **F-J:** mRNA levels of Proinflammatory cytokines *Il1β (*F), *Il6* (G), *Tnfα* (H), *Muc4* (I), and *Muc2* (J), analyzed by RT-qPCR, with Ct values normalized to *Actb* mRNA, in colon tissues of mice fed on RCD, MBD, MFD, and HFD after 7 days of DSS- or water-treatment (n = 8). Data are mean ± SEM of two or more indendent experiments, with statistical analysis by two-way ANOVA with Tukey’s multiple comparison test (B, C, D, F, G and H), ordinary one way ANOVA with RCD as reference (I, J). *p < 0.05, **p < 0.01, ***p < 0.005, ****p < 0.001

The disease severity score shows that both MBD and MFD-fed mice exhibit delayed onset of these disease symptoms, and the severity is significantly lower (Figure 1B). In contrast to both the RCD and HFD-fed mice, the reduction in body weight during the DSS challenge was also lower for MBD and MFD-fed mice (Figure 1C). Though DSS treatment led to a reduction in colon length in all groups, MBD and MFD-fed mice exhibited a lesser colon length reduction compared with both HFD and RCD after DSS treatment (Figure 1D & Figure S1C). Histopathology analysis showed that while all groups of mice exhibited immune cell infiltration in the lamina propria upon treatment with DSS, the RCD-fed and HFD-fed mice exhibited more immune cell infiltration and disintegration of the epithelial layer (Figure 1E). On the other hand, both MBD and MFD-fed mice exhibited well-maintained colonic epithelial architecture. Importantly, HFD-fed mice show immune cell infiltration even without DSS treatment, suggesting diet-induced inflammation (*24*).

DSS-treated mice fed with milk-based diets exhibited significantly lower mRNA levels of the pro-inflammatory cytokines *Il1b*, *Tnfa*, and *Il6* compared to those on HFD and RCD (Figure 1F-H), indicating a reduced inflammatory response in the milk diet groups. In contrast, the expression of the anti-inflammatory cytokines *Il10* and *Tgfb* did not change between RCD and both milk-based diet groups upon DSS treatment (Figure S1D&E), suggesting that the protective effect of milk diets stems from reduced pro-inflammatory signalling rather than enhanced anti-inflammatory activity. Notably, the HFD group showed reduced *Il10* and *Tgfb* expression following DSS administration (Figure S1D&E). Similarly, we observed higher levels of mucin gene mRNAs *Muc2* and *Muc4*, which are known to enhance the barrier function, in the milk-based diet-fed groups treated with DSS compared with the RCD group treated with DSS Figure 1I&J) (*25*, *26*).

DSS-induced colitis primarily functions by increasing gut permeability (*27*). To address whether the protection is specific to this mode of colitis induction, we used another approach, 2,4,6-trinitrobenzenesulfonic acid (TNBS), a hapten that induces transmural inflammation by activating the Th1 response (*28*). Interestingly, MFD did not show any protection against the TNBS model of colitis (Figure S1F-J). Interestingly, we also found that MFD failed to provide any protective effect when subjected to a much harsher (2.5%) treatment with DSS (Figure S1K-O). These results are consistent with the hypothesis that milk-based diets primarily enhance the gut barrier function, which can withstand mild DSS insults but fails if the barrier is breached with a higher concentration of DSS.

### Milk-based diets maintain gut barrier function after 1.5% DSS treatment

To understand the protective correlates in epithelial cells subjected to milk-based diets, we performed a single-cell RNAseq analysis of mice fed on MBD or RCD after 7 days of DSS treatment and compared them to untreated controls (Figure S2A). For each condition, cells from three mice were pooled. We sorted epithelial (EpCAM+), B cell (CD45+ CD19+), T cell (CD45+ CD3+), and non-lymphoid (CD45+ CD3− CD19−) cells from the colon (Figure S2B) and mixed (6000 epithelial cells, 2000 B cells, 4000 T cells, and 8000 myeloid cells) before subjecting them to GEM generation and library preparation using 10X genomics 3 GEX technology. The scRNA-Seq data were analysed primarily using the Seurat package (Figure S2C). Epithelial cells (*EpCAM*, *Krt8*, and *Krt18*), B cells (*CD79a*, *Ms4a1*, and *Cd19*), T cells (*Cd3d*, *Cd3e*, and *Cd3g*), and myeloid cells (*Cd14*, *Itgam*, *Fcgr3*, *Itgax*, *H2-Aa*, and *H2-Ab1*) were separated and analyzed (Figure S2C&D).

Re-clustering of the epithelial cells (Figures S2C & D) resulted in 23 distinct clusters (Figure 2A). Marker analysis identified all major epithelial cell types within these clusters (Figure 2B and Figures S3A–G). The control groups from both diets exhibited more or less similar distributions of epithelial cell types (Figure 2B and Figure S3H). Various studies have shown that colitis induction in mice and LPS stimulation in epithelial cells lead to increased expression of *Il1b*, *Tnfa*, and *Cd14* in colonocytes (*29*, *30*). In line with this, DSS-treated RCD-fed mice showed a dramatic expansion of clusters 0, 3, 8, 9, 15, 19, 20, 21, and 22 (Figure 2A&B and Figure S3H), which express high levels of inflammatory genes, including *Il1b* & *Tnfa* (Figure 2C and Figure S3I). We therefore classified these clusters as inflamed epithelial cells (Figure 2B). Interestingly, these cells are almost exclusively present in the RCD-DSS group, with very low numbers in the MBD-DSS group (Figure 2B and Figures S3H&J), reiterating the reduced inflammation in the latter group.

**Figure 2:**
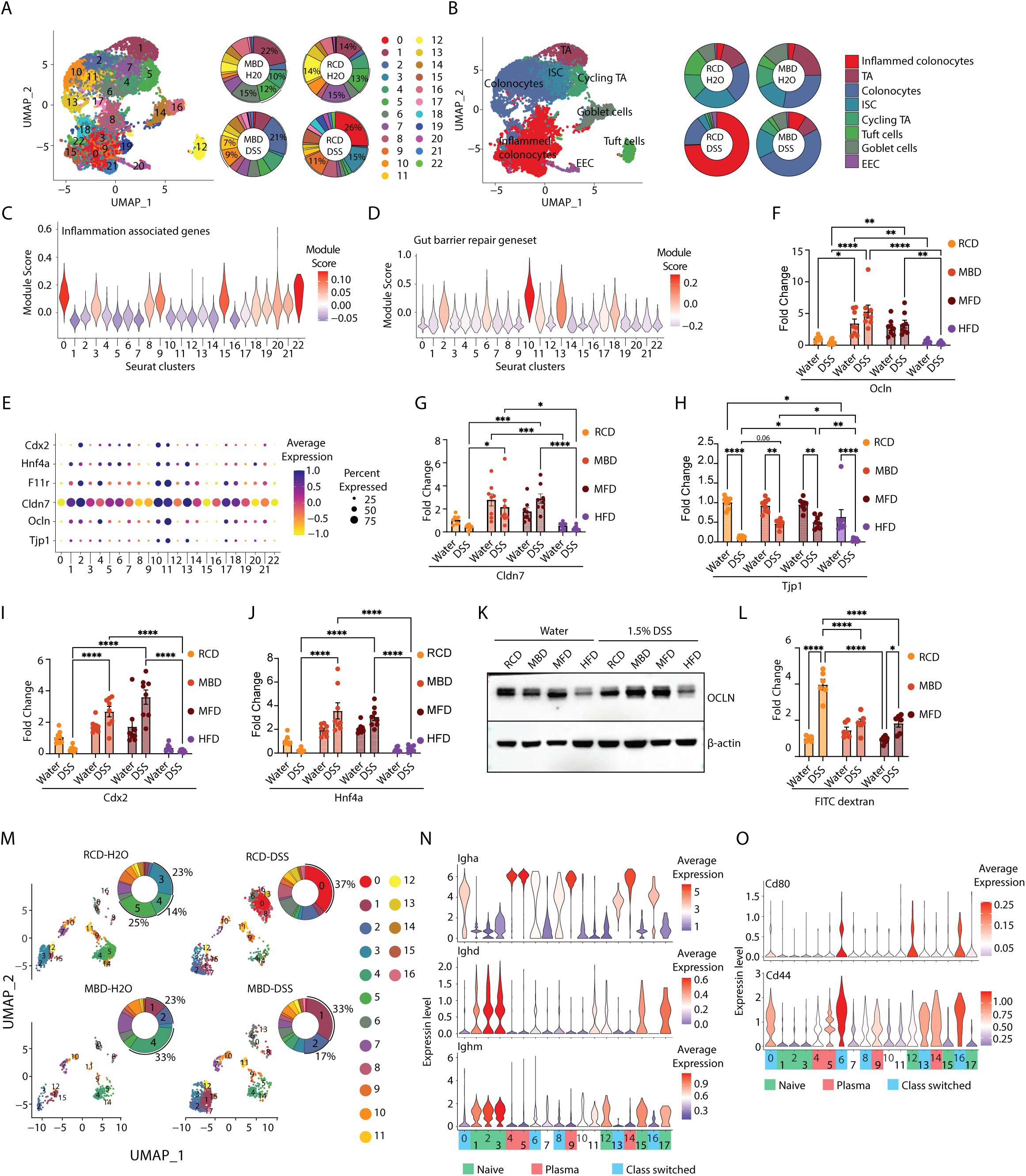
Milk-based diets maintain gut barrier function even after DSS treatment. **A:** UMAP and donut plots showing the distribution of epithelial cells from different experimental groups across different clusters. The experimental groups are RCD-H2O (RCD on water), RCD-DSS (RCD on 1.5% DSS), MBD-H2O (MBD on water), and MBD-DSS (MBD on 1.5% DSS). The percentage of cells belonging to indicated clusters is shown next to each donut plot. **B:** UMAP and donut plots showing the distribution of epithelial cells from different experimental groups across different cell types. **C&D:** Violin plot showing module scores for inflammatory (C) and gut barrier repair associated gene (D) sets across all clusters (Table S2). **E:** Dot plot showing expression of gut barrier-associated genes across clusters. The color indicates the expression level, and the size of the dots indicates the percentage of cells expressing the gene. **F-H:** Expression of gut barrier-associated genes *Ocln* (F), *Cldn7* (G), and *Tjp1* (H) analyzed by RT-qPCR, with Ct values normalized to *Actb*, in colon tissues of DSS- or water-treated mice fed RCD, MBD, MFD, and HFD (n= 8). **I&J:** Expression levels of transcription factor genes, *Cdx2* (I) and *Hnf4α* (J), analyzed by RT-qPCR. Ct values were normalized to *Actb*, in colon tissues of DSS- or water-treated mice fed RCD, MBD, MFD, and HFD (n= 8). **K:** Western blot analysis of distal colon tissue isolated from mice fed on RCD, MBD, MFD, and HFD with and without DSS treatment, showing OCLN. **L:** Gut permeability assay, measuring fluorescence in the serum of DSS- or water-treated mice fed RCD, MBD, MFD, and HFD 6 hours after oral gavage of FITC-dextran (n= 6). **M:** UMAP and donut plots showing the distribution of B cells from different experimental groups across different clusters. The experimental groups are RCD-H2O (RCD on water), RCD-DSS (RCD on 1.5% DSS), MBD-H2O (MBD on water), and MBD-DSS (MBD on 1.5% DSS). B cells were identified based on markers *Cd79a, Ms4a1,* and *Cd19*. The percentage of cells belonging to some of the clusters is shown next to each donut plot. **N:** Violin plot for average expression of I*gha, Ighd,* and *Ighm* across the B cell clusters to show naive, plasma, and class-switched B cells. Cell types are color coded on the X axis. **O:** Violin plot for average gene expression for *Cd80* and *Cd44*, markers for activated B cells. Data are mean ± SEM of two independent experiments, with statistical analysis by two-way ANOVA with Tukey’s multiple comparison test (F, G, H, I, J and L). *p < 0.05, **p < 0.01, ***p < 0.005, ****p < 0.001.

On the other hand, the MBD-DSS group exhibited an increased number of colonocytes and preservation of the ISC population compared with the RCD-DSS group (Figure 2B, Figure S3J). Notably, clusters 2, 10, and 11, classified as colonocytes, were upregulated in the MBD-DSS group compared with the RCD-DSS group (Figure 2A and Figure S3H). A meta-analysis of transcriptomics data from colitic and normal colonic tissue identified a set of 10 marker genes for mucosal barrier repair (*31*). We found that clusters 2, 10, 11, 13, and 18 exhibited increased expression of this gene set, with the highest expression in cluster 10, which was predominantly enriched in the MBD-DSS group compared with RCD-DSS group (Figure 2D).

Further analysis of the expression of gut barrier junction protein-coding genes showed upregulation of *Tjp1*, *Ocln*, *Cldn7*, and *F11r* in these clusters (Figure 2E). These clusters also showed elevated expression of transcription factors *Hnf4a* and *Cdx2*, both of which have been previously associated with gut barrier integrity (*32–35*). To validate these findings, we performed qRT-PCR analysis on colon tissues from control and DSS-treated mice (Figure 2F-J). Expression levels of *Tjp1*, *Cldn7*, and *Ocln* mRNAs were significantly upregulated in the MBD and MFD-fed mice treated with DSS, compared to those in the RCD-DSS group. Additionally, *Cdx2* and *Hnf4a* mRNA levels recapitulate the scRNA-seq results. Interestingly, these genes were expressed at lower levels in the HFD group, further supporting their association with gut barrier protection and repair. Further, protein levels of Occludin also show higher levels in DSS-treated, MBD- and MFD-fed mice compared with their RCD and HFD counterparts, confirming upregulation of tight junction proteins (Figure 2K). To assess whether the increased expression of tight junction proteins translates into improved barrier function, we measured intestinal permeability. Indeed, we observed significantly lower levels of FITC-dextran in the blood 6 hours after oral gavage in both MBD and MFD-fed mice (Figure 2L). These results clearly indicate that milk-based diets maintain the colonic epithelial barrier function even after a 1.5% DSS challenge.

### MBD-fed mice exhibit muffled Immune activation after DSS treatment

Next, we examined the effects of MBD on the DSS treatment-associated immune activation in the colon. (Figure S2B-D). B cell analysis revealed 17 clusters grouped into three major categories: naïve (*IgD*⁺*/IgM*⁺, clusters 1, 2, 3, 12, 15, 17), plasma (clusters 4, 5, 9, 14), and class-switched *IgA*⁺ B cells (clusters 0, 6, 8, 13, 16) (Figure S4A). While *IgA* expression was highest in plasma cells, class-switched clusters showed moderate *IgA* along with elevated activation markers *CD80* and *CD44* (Figures 2M-O). Control mice on both diets showed comparable levels of plasma and naïve B cells, though with distinct cluster distributions: RCD controls had more cells in cluster 3, while MBD controls favoured clusters 1 and 2 (Figure 2M). Upon DSS treatment, these MBD clusters expanded further. Strikingly, the MBD-DSS group was dominated by naïve B cells, particularly cluster 1 (33%), whereas the RCD-DSS group was enriched in class-switched *IgA*⁺ B cells, with cluster 0 accounting for 37% of total B cells (Figure 2M-O and S4A). These results suggest that milk-based diets reduce antigen exposure of B cells, as evidenced by the naïve B cell enrichment and lower IgA^+^ antigen-experienced B cells in MBD-DSS mice compared to the activated B cell profile in RCD-DSS mice. This could be the result of a stronger epithelial barrier function.

T cells clustered into 17 groups, with 62% of cells distributed across clusters 0–4 (Figure S4B). Clusters 0 and 1 expressed *Cd8a* without *Cd8b1*, likely representing gut-resident CD8αα cells. Clusters 2, 3, 9, 10, 12, 13, 14, and 16 represented CD8⁺ T cells co-expressing *Cd8a* and *Cd8b1* (Figure S4C&D). In contrast, clusters 4, 5, 7, 8, 11, and 15 were CD4⁺ T cells, which included distinct subsets: Th1 (*Cd4, Tbx21, Ifng*; cluster 4), Th2 (*Gata3, Il13, Il4*; cluster 11), Tregs (*Foxp3, Il2r*; cluster 7), and Th17 (*Rorc, Il17*; cluster 15) (Figure S4C&D). Unlike B cells, T cell distributions across dietary and treatment conditions were generally similar. The only notable difference was an enrichment of activated CD8⁺ T cells (cluster 2) in chow-fed control mice. In DSS-treated groups, we observed a modest increase in Th1 (cluster 4) and Treg (cluster 7) populations in MBD-fed mice (Figure S4E).

Analysis of myeloid cells revealed no major differences between MBD and RCD fed mice under non-inflammatory conditions (Figure S4F&G). Both groups showed enrichment of gut-protective macrophages (*Mrc1/CD206*, *Csf1r*; cluster 2), *Ly6c2*-expressing monocytes (cluster 5), and MHCII⁺ cDC2 cells (cluster 8) (Figures S4G–J). Upon DSS treatment, notable shifts were observed. RCD-fed mice showed an expansion of cytotoxic NK cells (cluster 0), marked by *Cd27*, *Klrk1*, *Gzma*, *Gzmb*, *Prf1*, *Fasl*, and *Trail* (Figures S4G&K). These cells also exhibited elevated ribosomal gene expression, indicating increased biosynthetic activity (Figure S4L). In contrast, the MBD-DSS group preserved macrophages and showed an increase in inflammatory monocytes (cluster 3), expressing *Il1a*, *Il1b*, *Saa3*, *Vim*, and *Fn1* (Figure S4M). Together, these findings suggest that DSS-treated RCD-fed mice develop a robust cytotoxic NK cell response, while MBD-fed mice maintain inflammatory monocytes and homeostatic macrophages.

Taken together, these results indicate that MBD-fed mice exhibit a distinct immunological profile, characterised by reduced levels of antigen-experienced B cells and cytotoxic NK cells, along with increased levels of homeostatic macrophages collectively reflecting lower immune activation.

### Milk-based diets elicit protective effects against gut inflammation by activating the JNK2 pathway in the epithelial cells

Next, we investigated the mechanisms by which milk-based diets affect intestinal barrier function. We performed the regulome analysis of the scRNAseq data using the SCENIC package (*36*). We observed that the clusters that are enriched in the MBD groups 2, 10, and 11 showed enrichment of a common set of regulome (Figure 3A and Figure S5A). The transcription factors whose regulome is enriched in these clusters include several AP-1 transcription factors (*Jund*, *Jun*, *Junb*, *Fosl*, *Atf3*, and *Atf4*) and transcription factors well established to have positive effects on gut barrier function (*Hnf4a*, *Nr1i2*/PXR, *Nr1h4*/FXR, *Egr1*) (*37–39*). Transcript levels of each of these transcription factors are also upregulated in these clusters (Figure 3B). qRT-PCR analysis on IECs from non-DSS-treated mice confirmed increased expression of *Jun*, *Junb*, and *Atf3* in the milk-based diet groups, but not in the HFD group (Figure S5B)

**Figure 3:**
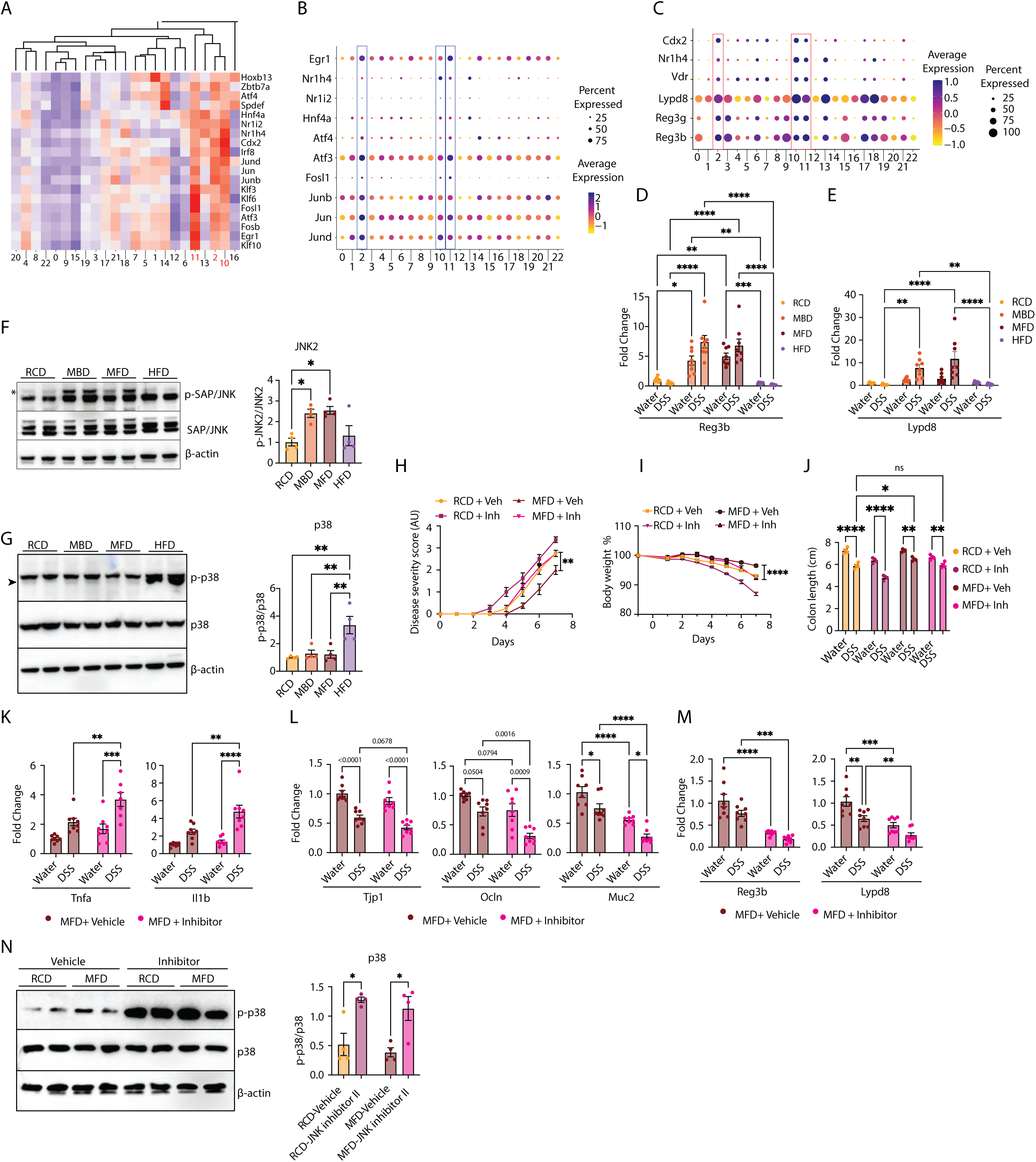
Milk-based diets elicit protective effects against gut inflammation by activating the JNK2 pathway in the epithelial cells. **A:** Heatmap showing the regulome expression calculated using the SCENIC package, highlighting transcription factors whose targets are enriched in clusters 2, 10, and 11. **B:** Dot plot showing expression levels of gut barrier-associated transcription factors identified in SCENIC analysis and enriched in clusters 2, 10, and 11. **C:** Dot plots of antimicrobial protein (AMP) genes (*Reg3b, Reg3g,* and *Lypd8*) and associated transcription factors (*Vdr, Cdx2,* and *Nr1h4*) across epithelial cell clusters, highlighting the enrichment of these genes in clusters 2, 10, and 11. **D&E:** Expression levels of AMP genes *Reg3b* (D), and *Lypd8* (E), and analyzed by RT-qPCR, with Ct values normalized to *Actb*, in colon tissues of DSS- or water-treated mice fed RCD, MBD, MFD, and HFD (n= 8). **F&G:** Western blot analysis of intestinal epithelial cells (IECs) isolated from colons of mice subjected to RCD, MBD, MFD, and HFD (without DSS treatment), probed with antibodies against p-JNK, JNK, p-p38, and p38 (n= 4). Representative blot (left panel) and densitometric quantification (right panel) are shown. In the JNK blot, the upper band (indicated by an asterisk) shows JNK2, while the lower band shows JNK1. In the P38 blot, the lower band (indicated by the arrowhead) indicates the P38; the upper band is a non-specific band. **H-J:** JNK inhibitor II or vehicle was administered to RCD and MFD-fed mice 24 hours before treatment with DSS. Disease severity score (H), body weight percentage (I), and total colon length in cm(J) of DSS and water groups are plotted (n= 4). **K-M:** mRNA levels of proinflammatory cytokines (*Tnfα* and *Il1b*) (K), gut barrier genes (*Tjp1, Ocln,* and *Muc2*) (L), and AMPs (*Reg3b* and *Lypd8*) (M), in the colon tissue of JNK inhibitor II treated mice were analyzed by RT-qPCR, with Ct values normalized against *Actb* (n= 8). Dietary groups are indicated. **N:** p-p38 and p38 levels were analyzed in IECs from mice fed on RCD and MFD, isolated 7 days after p-JNK inhibition (n= 4). Representative blot (left panel) and densitometric quantification (right panel) are shown. Data are mean ± SEM of two independent experiments, with statistical analysis by two-way ANOVA with Tukey’s multiple comparison test (D, E, H, I, J, K, L, M, and N) and one-way ANOVA with Tukey’s multiple comparison test (F, and G). *p < 0.05, **p < 0.01, ***p < 0.005, ****p < 0.001.

AP-1 transcription factors are key regulators of inflammation and epithelial regeneration. Constitutive activation of AP-1 is known to promote intestinal stem cell proliferation (*16*). Additionally, AP-1 functions as a central hub within a transcriptional network that modulates the expression of tight junction proteins and antimicrobial peptides (Figure S5C)(*37*, *38*, *40–44*). Consistent with this, our data showed upregulation of transcripts encoding tight junction proteins in the milk-diet groups (Figure 2F–H). Furthermore, transcripts of antimicrobial proteins such as *Reg3b* and *Lypd8,* known to support host-beneficial microbiome composition, were highly expressed in clusters 2, 10, and 11 (Figure 3C). qPCR analysis of whole colon tissues confirmed increased expression of these genes in the milk-based diet groups (Figures 3D&E), but not in the HFD group. In contrast, the HFD group showed elevated expression of *Cox2* and *Ptges2* mRNAs, which are involved in prostaglandin synthesis and contribute to inflammation (Figure S5D).

MAPK pathways such as ERK, JNK, and P38, along with several GPCR pathways, are known to activate AP-1 transcription factors. To determine whether any of these pathways are modulated by milk-based diets, we performed western blot analysis on colonic epithelial cells isolated from control mice on different diets (without DSS treatment). Interestingly, milk-based diets induced phosphorylation of JNK2 at Thr183/Tyr185 (Figure 3F, asterisk), indicating that these epithelial cells are primed for JNK2 activation even in the absence of inflammatory stimuli. In contrast, mice on a high-fat diet (HFD), which is associated with increased baseline gut inflammation, showed elevated phosphorylation of p38 (Thr180/Tyr182) instead of JNK2 (Figure 3G, arrowhead). These findings suggest that milk-based and lard-based high-fat diets activate distinct MAPK pathways in colonic epithelial cells under non-inflammatory conditions. We hypothesise that milk-based diets prime the gut epithelium by selectively activating the JNK2 pathway in epithelial cells, potentially contributing to enhanced barrier protection and reduced inflammatory responsiveness, while HFD activates P38 pathway, which is associated with gut inflammation (*20–22*, *45*).

### Inhibition of the JNK2 pathway abolishes the protective effect

Deletion of JNK2 has been shown to exacerbate DSS-induced colitis, even at mild doses of DSS (*46*). JNK2 signalling in epithelial cells is critical for the maintenance of epithelial integrity during inflammatory injury (*18*). So, we investigated whether JNK2 activation mediates the protective effects of milk-based diets. To test this, we rectally administered a JNK inhibitor (SP600125; JNK inhibitor II) at a dose of 10 µg/g body weight. This treatment effectively reduced JNK2 activity in the colonic epithelium within 24 hours (Figure S5E). We then treated MFD-fed mice with the JNK inhibitor 24 hours before DSS exposure. Seven days after DSS treatment, inhibitor-treated MFD-fed mice displayed increased disease severity, greater body weight loss, and shorter colon length compared to vehicle-treated controls (Figures 3H–J, S5F). Notably, JNK inhibition also worsened disease outcomes in RCD-fed mice, suggesting that even basal levels of JNK activation in this group contribute to gut protection. Histopathological analysis further confirmed increased tissue damage in the MFD group following JNK inhibition (Figure S5G). Consistent with this, *Tnf*α and *Il1*β mRNA levels were significantly upregulated in the JNK-inhibited groups, indicating elevated inflammation (Figure 3K). In parallel, the expression of barrier protection-associated genes *Muc2*, *Tjp1*, *Ocln*, and *Hnf4*α was reduced, suggesting that JNK positively regulates these genes (Figure 3L, Figure S5H). Additionally, antimicrobial peptide genes *Lypd8* and *Reg3b* were also downregulated upon JNK inhibition (Figure 3M). Notably, JNK2 inhibition persisted even after 7 days in MFD-fed mice without DSS treatment (Figure S5I). This sustained inhibition suggests that once JNK2 is suppressed, dietary intervention is insufficient to restore its activity. Interestingly, JNK2 inhibition was associated with increased phosphorylation of p38, aligning with the heightened disease severity observed in the JNK2 inhibitor-treated groups (Figure 3N). These findings highlight the opposing roles of the JNK2 and p38 pathways in regulating colonic inflammation.

The JNK pathway plays a critical role in regulating epithelial proliferation, barrier function, apoptosis, and enterocyte survival under stress conditions (*15*, *18*, *47*, *48*). We hypothesised that enhanced epithelial renewal may contribute to the protective effects of milk-based diets. However, western blot analysis of Cyclin D1 levels and microscopic analysis of Ki67 positive proliferative zones showed that neither of the milk-based diet groups exhibited any differences in these two markers of proliferation (Figure S5J&K). However, Ki67 staining of colon tissues revealed a marked reduction in the proliferative zone in HFD-fed mice, suggesting impaired epithelial renewal in this group. The mRNA levels of apoptotic markers Bax, Puma, and Noxa and the levels of cleaved Caspase-3 were also unchanged in the milk-based diet groups. HFD-fed mice, on the other hand, exhibited increased levels of these markers, indicating elevated apoptosis (Figure S5L&M). Together, these findings suggest that while HFD-fed mice exhibit defects in epithelial proliferation and increased apoptosis, which may contribute to impaired intestinal homeostasis, milk-based diet-fed mice did not exhibit any changes in proliferation and apoptosis markers.

We further examined the upstream activators of JNK2 and P38, MKK4 and MKK7, by performing western blot analysis on IECs of different diet groups. We observed that MKK7, the primary activator of JNK2, showed no change across the different diet groups (Figure S5N) (*49*). However, MKK4, which can also phosphorylate p38, was significantly elevated in the HFD-fed group (Figure S5O) (*50*). These results suggest that HFD feeding activates P38 through MKK4 activation. However, how MFD activates JNK2 is unclear.

### *Dubosiella Newyorkensis*, enriched in the milk-based diet groups, provides gut protection

Milk components are known to affect the gut microbiome (*51*, *52*). The changes in the expression levels of several antimicrobial peptides in mice fed on milk-based diets also point toward the modulation of the gut microbiome by milk-based diets. To test whether any of the protective correlates of the milk-based diets are due to the gut microbiome, we removed the microbiota from these mice by treatment with a cocktail of antibiotics (Vancomycin (0.5 mg/ml), Ampicillin (1 mg/ml), Neomycin (0.5 mg/ml), and Metronidazole (0.625 mg/ml) in 2% sucrose) (Figure 4A & Figure S6A). When milk-based diet-fed mice were treated with antibiotics, the epithelial JNK2 phosphorylation was reduced significantly compared with MFD without antibiotics (Figure 4B). Interestingly, the genes that were associated with protection, such as the transcription factor *Hnf4*α and the AMPs *Reg3b* and *Lypd8,* and tight junction proteins were downregulated compared to those of the RCD group in the MFD-ΑΒΧ group (Figure S6B-D), suggesting that the microbiome plays an important role in the expression of protective markers in milk-based diet groups.

**Figure 4:**
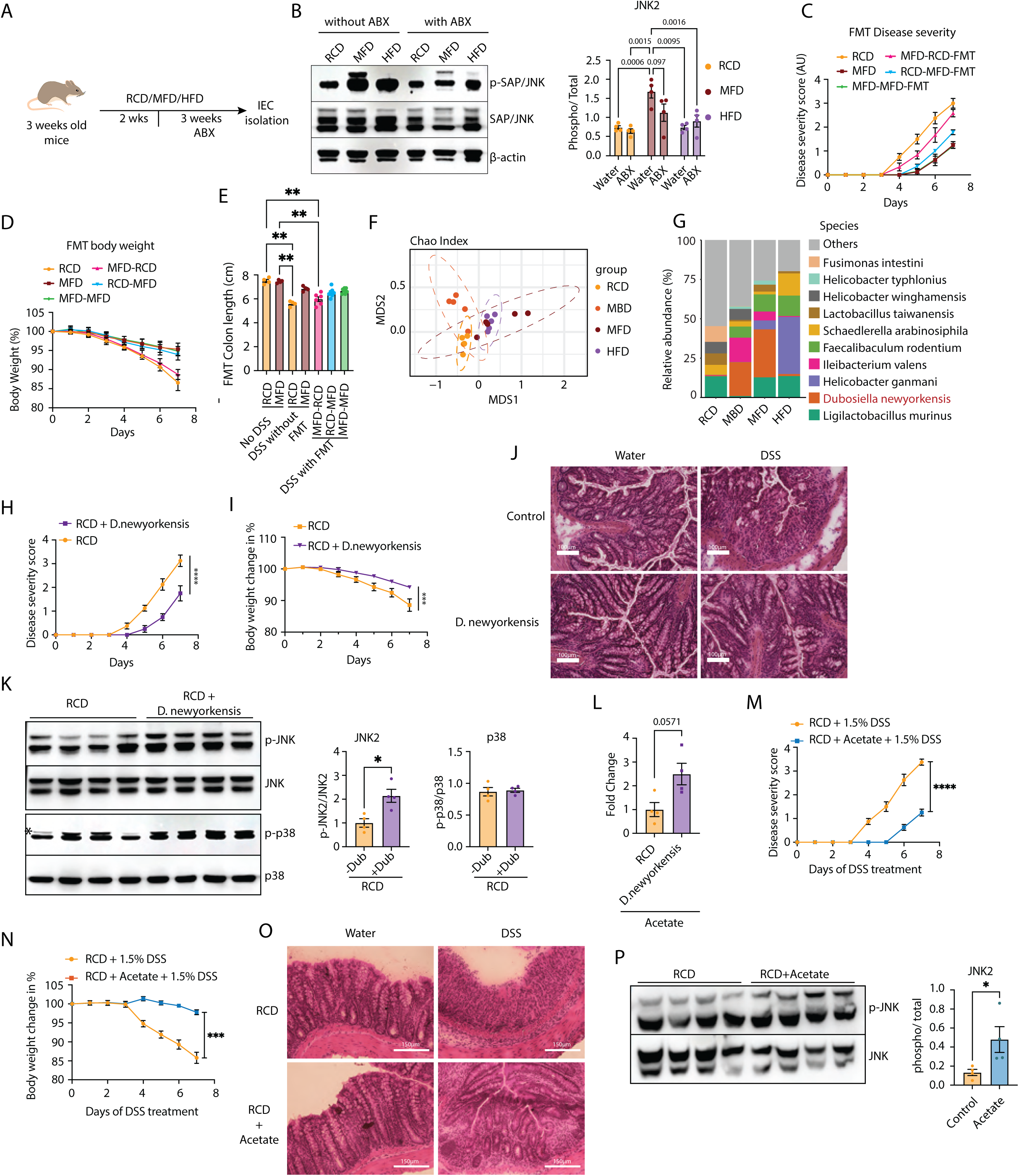
Dubosiella Newyorkensis, enriched in the milk-based diet groups, provides gut protection via Acetate. **A:** Schematic illustration of the experimental workflow for antibiotic treatment. **B:** Western blots of IECs isolated from mice with or without antibiotic (ABX) treatment, fed RCD, MFD, and HFD (n= 4), showing levels of p-JNK and total JNK (left panel). Quantification of the blots is on the right panel. **C-E:** FMT from RCD to MFD (MFD-RCD-FMT) (n = 6), MFD to RCD (RCD-MFD-FMT) (n = 7), MFD to MFD (MFD-MFD-FMT) (n = 7), control RCD andMFD (n =4) are shown. Disease severity score (C), body weight percentage (D), and total colon length in cm (I) in DSS and water groups are plotted. **F:** Microbial beta-diversity index, Chao index of fecal microbiome isolated from mice fed on RCD, MBD, MFD, and HFD (n = 5). **G:** Stacked bar plot showing the abundance of the top 10 species across different experimental groups. D. newyorkensis is highlighted. **H&I:** Disease severity score (H), and body weight (I) of control and D.newyorkensis treated mice challenged with DSS (n = 4). J: Representative images of H&E-stained colon tissue sections of control and D.newyorkensis treated mice challenged with DSS. (Scale bars: 100 μm). **K:** Western blot analyses of IECs isolated from mice treated with D. newyorkensis, showing p-JNK, total-JNK, p-p38 and total p38 levels (left panel) (n = 4). Quantification of the blots is on the right panel. **L:** Levels of acetate in the fecal matter of mice treated with D. newyorkensis and control mice. **M&N:** Disease severity score (M), and body weight (N) of acetate and DSS-treated mice (n = 4). **O:** Representative images of H&E-stained colon tissue sections of control and acetate treated mice challenged with DSS. (Scale bars: 150 μm). **P:** Western blots of IECs isolated from mice with or without acetate treatment, showing levels of p-JNK and total JNK (n = 4)(left panel). Quantification of the blots is on the right panel. Data are mean ± SEM of two independet experiments, with statistical analysis by two-way ANOVA with Tukey’s multiple comparison test (B, C, D, I, and N), two-way ANOVA with Šídák’s multiple comparison test (H, and M), Kruskal-wallis test with Dunn’s multiple comparison test (E), and Mann Whitney test (K, L, and P). *p < 0.05, **p < 0.01, ***p < 0.005, ****p < 0.001.

Similarly, the P38 activation was also reduced in HFD-fed mice when treated with antibiotics (Figure S6E). mRNA levels of genes downstream to P38, such as *Cox2* and *Ptges2*, and *Tnf*α, a pro-inflammatory cytokine, were also reduced in the HFD group treated with antibiotics (Figure S6F). In addition, the lower levels of tight junction protein and mucin expression observed in HFD-fed mice were restored by antibiotic treatment (Figure S6D). This supports the previous observations that in HFD-fed mice, an altered microbiome is the primary inducer of inflammation (*53–55*). These results indicate that, as in the case of the hyperinflammation induced by HFD, the protective effect of MFD is also dependent on the microbiome.

To further investigate the role of the microbiome in the protective effect, we performed faecal microbiota transplantation (FMT). The gut microbiota of recipient mice fed either MFD or RCD was depleted using PEG 6000 (Figure S6G), followed by faecal matter transplantation from donor groups (*56*), for three weeks. This was followed by treatment with 1.5% DSS. We included five groups, each comprising seven mice: (1) RCD recipients with MFD donors (RCD-MFD-FMT), (2) MFD recipients with RCD donors (MFD-RCD-FMT), (3) MFD recipients with MFD donors (MFD-MFD-FMT), (4) MFD control, and (5) RCD control. Interestingly, both RCD-MFD-FMT and MFD-MFD-FMT groups exhibited protective effects, whereas the MFD-RCD-FMT group showed significant weight loss, reduced colon length, increased disease severity and severe epithelial damage (Figure 4C-E and Figure S6H&I). Taken together, these findings clearly indicate that the microbiome plays a crucial role in mediating the protective correlates of milk-based diets.

Next, we analysed the composition of the gut microbiome by 16s rRNA sequencing using the Oxford Nanopore platform, which allows reliable species-level taxonomic identification by sequencing the full-length 16s loci (*57*). Interestingly, even though both MFD and MBD showed protection against the DSS-induced pathology, they exhibited little similarity in terms of microbial composition. MBD showed the highest alpha diversity among the groups, while both MFD and HFD groups exhibited lower alpha diversity, suggesting that the overall diversity is affected mostly by the macronutrient content of the diets (Figure S7A&B). Furthermore, the Chao index also indicated that bacterial composition among the MFD and MBD is divergent (Figure 4F). Differentially enriched microbes in different diet groups show that both MBD and MFD have increased protective species in the microbiome, while HFD shows increased levels of known pathogenic bacteria (Supplementary Dataset 1).

Despite the difference between MBD and MFD in the overall composition of the microbiome, we observed that *Dubosiella newyorkensis* is one of the most abundant species in both groups (Figure 4G and Figure S7C-E). *D. neworkensis*, which is a mouse homolog of the human symbiont *Clostridium innocuum*, has been recently shown to elicit gut protective properties (*58*). To address whether *D. newyorkensis* indeed mediates the protective effects of MBD and MFD, we subjected the mice to oral treatment with *D. Newyorkensis* for 2 weeks, followed by DSS treatment (Figure S7F). They exhibited protection from DSS-induced inflammation as evidenced by reduced disease severity score, maintenance of body weight, and tissue damage (Figure 4H-J). These data clearly show that *D. newyorkensis* protects mice from DSS-induced colitis.

We next checked the protective correlates we observed in the milk-based diet groups (Figure S7G-J). The mRNA levels of pro-inflammatory cytokines TNFα and IL1β were reduced in the *D. newyorkensis* group (Figure S7G), while the transcripts for the barrier-protective proteins CLDN7, OCLN, TJP1, MUC2, and HNF4α were upregulated as seen in milk-based diet groups (Figure S7H). Apart from the low inflammatory genes and high gut barrier protection-associated genes, the transcript levels of anti-microbial protein transcripts *Reg3b* and *Lypd8* were upregulated in the *D. newyorkensis* treated group (Figure S7I). These results clearly show that *D. newyorkensis* recapitulates salient features of milk-based diets in protection against gut inflammation. Moreover, western blot analyses of the epithelial cells show that phospho-JNK2 is indeed upregulated in the *D. newyorkensis* treated group (Figure 4K). On the other hand, we did not observe any difference in the p-P38 levels, suggesting that *D. newyorkensis* is sufficient to selectively upregulate the JNK2 pathway in the colonic epithelium (Figure 4K) and enable the protection from DSS-induced inflammation.

### Acetate induces the JNK2 pathway in the colonic epithelium to enhance gut barrier function

Short-chain fatty acids are major contributors to gut barrier integrity (*59*, *60*). The commensal bacteria produce short-chain fatty acids (SCFAs) after fermenting the dietary fibre. Moreover, *D. newyorkensis, which* is enriched in milk-based diets, also produces short-chain fatty acids (*58*). We performed GC-MS analysis of faecal samples of mice subjected to RCD, MFD, and HFD to quantify different short-chain fatty acids. Compared with RCD and HFD fed mice, we found that acetate, propionate, and butyrate are high in the faecal samples in MFD-fed mice (Figure S8A). On the other hand, in HFD-fed mice, all three of these short-chain fatty acids were low, clearly showing that these short-chain fatty acid levels are indeed correlated with gut protection against DSS treatment. Interestingly, faecal samples of mice treated with *D. newyorkensis* also exhibited higher levels of acetate compared with control samples (Figure 4L). On the other hand, propionate and butyrate did not show a statistically significant increase in these mice (Figure S8B). This data suggests that acetate could be the driving force behind the protective phenotype.

Acetate has been shown to protect against DSS-induced colitis previously; however, the mechanism is not clearly understood (*61*, *62*). To test if acetate treatment functions through JNK2 phosphorylation, we provided potassium acetate to these mice in drinking water (200 mM). Indeed, we found that acetate treatment provided protection against DSS-induced pathologies based on body weight, colon length, disease severity score, and tissue destruction (Figure 4M-O & Figure S8C&D). The mRNA levels of pro-inflammatory cytokines *Tnf*α, *Il1*β, and *Ifn*γ were also significantly downregulated in the acetate-treated group, suggesting reduced inflammation (Figure S8E). Moreover, we observed increased mRNA levels of tight junction protein *Ocln*, mucin gene *Muc2*, and the transcription factor *Hnf4*α, which were all associated with MFD feeding in these mice (Figure S8F). Next, we asked if acetate enhances protection through the JNK2 pathway, as observed for milk-based diets and *D. newyorkensis*. Indeed, acetate treatment was sufficient to induce JNK2 phosphorylation in the epithelium, while it did not show any statistically significant effect on P38 phosphorylation (Figure 4P & Figure S8G). Similarly, the mRNA levels of AMPs *Reg3b* and *Lypd8* were also upregulated in the acetate-treated group (Figure S8H).

Acetate is known to modulate gut homeostasis either at the chromatin level, through histone acetylation, or via signalling pathways, primarily through the GPR43/FFAR2 receptor (*63*). GPR43, present in the colonic epithelium, senses SCFAs and modulates gut homeostasis, thereby providing protection against DSS-induced colitis (*64*). To investigate whether acetate affects JNK2 phosphorylation in colonic epithelial cells via GPR43 signalling, we rectally administered the GPR43 inhibitor GLPG097 (10 µg/g) in two doses prior to IEC isolation. Western blot analysis revealed that phospho-JNK2 levels in the MFD group remained unchanged despite GPR43 inhibition (Figure S8I). These findings suggest that JNK2 phosphorylation is not directly regulated by the GPR43 signalling pathway. The role of chromatin modifications on JNK2 activation by acetate may involve chromatin-level regulation, which needs further investigation.

Taken together, these results show that post-weaning milk-feeding supports the expansion of the *D. newyorkensis* population to increase short-chain fatty acid production, which in turn activates the epithelial JNK2 pathway to enhance gut barrier function. On the other hand, P38 activation is associated with diets that induce chronic inflammation in the gut.

### Milk supplementation during the weaning period is essential to establish the gut-protective microbiome and JNK2 phosphorylation

Next, we asked if the milk-based diets could extend their protective effect against repeated DSS challenges. To address this, we subjected these mice to a second round of DSS treatment after recovering for two weeks from the first round (Figure 5A). Interestingly, the disease severity scores, body weight changes, colon length, and pro-inflammatory cytokine levels showed that mice weaned onto MFD exhibited reduced inflammation even on repeated DSS challenges (Figure 5B-E; MFD(I&II)). However, initiating the MFD feeding after the first round of DSS treatment of RCD-weaned mice failed to show a protective effect (Figure S9A-C; RCD(I)-MFD(II)), suggesting that though milk-based diets can provide long-term protection from DSS-induced inflammation, this protective effect is lost once regular chow feeding is initiated. These results and the inability of MFD to reactivate JNK2 after one round of inhibition (Figure S5I) suggested that the MFD feeding must start during weaning to maintain gut barrier function.

**Figure 5:**
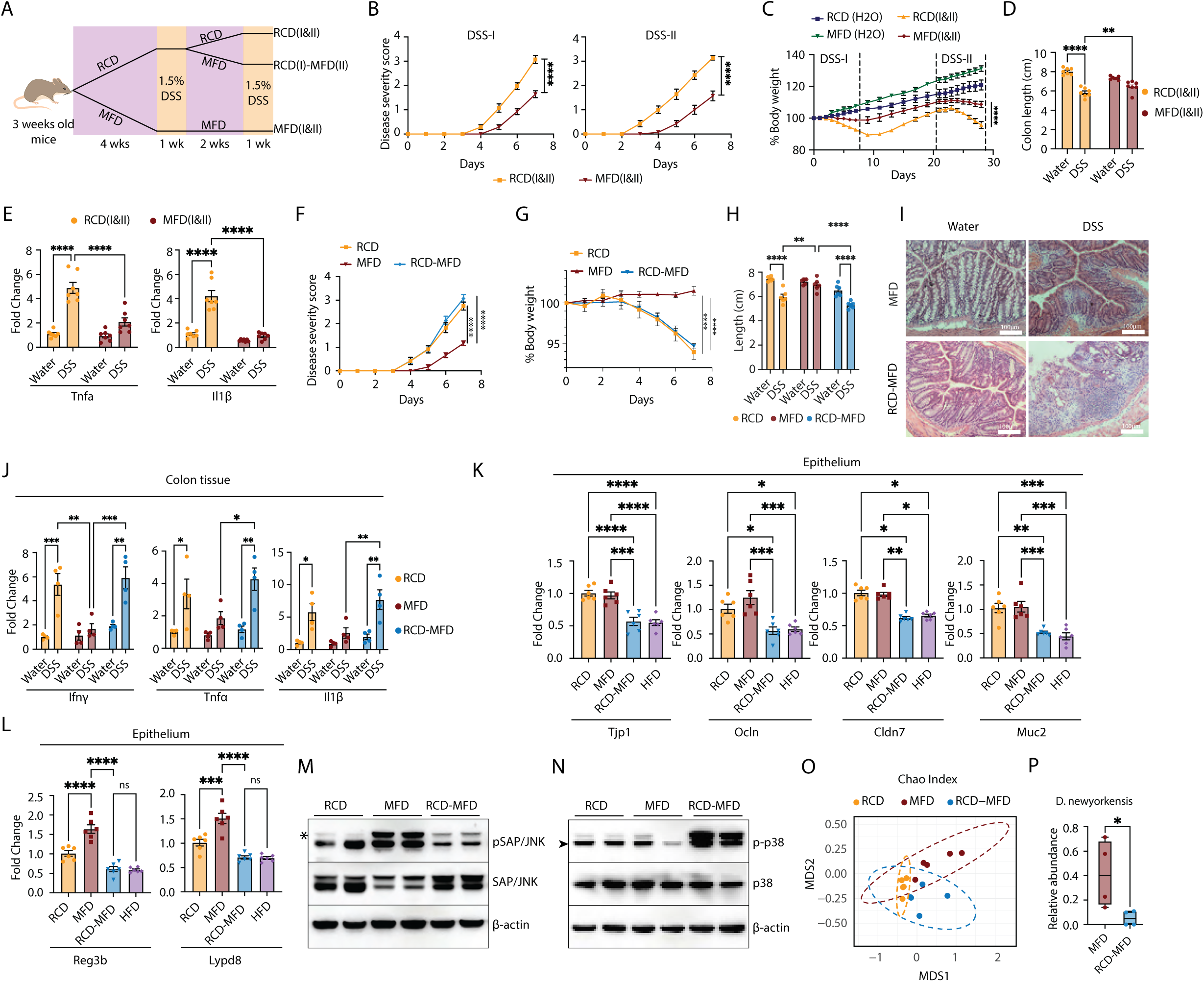
Milk supplementation during the weaning period is essential to establish the gut-protective microbiome and JNK2 phosphorylation. **A:** Schematic illustration of repeated DSS exposure experimental workflow. **B:** Disease severity score of RCD(I&II) and MFD(I&II) groups during the first DSS challenge (DSS-I) and second DSS challenge (DSS-II) (n = 7). **C:** Daily body weight (as the percentage with the day 0 as 100%) of mice in RCD(I&II) and MFD(I&II) groups along with their control groups (without any DSS treatment) (n = 7). The dashed vertical lines demarcate the DSS treatment periods (DSS-1 and DSS-II). **D:** Colon lengths of mice in cm of RCD(I&II) and MFD(I&II) groups, along with their control groups (without DSS treatment) (n= 7), were measured at the end of the experiment and plotted. **E:** Expression levels of proinflammatory cytokines (*Tnfα* and *Il1b* were analyzed by RT-qPCR, with Ct values normalized to *Actb*, in colon tissues of repeated DSS exposure and control groups (n = 7). **F-H:** Disease severity scores (F), body weight change (G), and total colon length in cm (H) of mice treated with 1.5% DSS and subjected to RCD, MFD, and RCD-MFD diets (n = 6). **I:** Representative images of H&E stained colon tissues from DSS- or water-treated mice fed on either MFD or RCD-MFD diets. (Scale bars: 100 μm). **J:** Expression levels of proinflammatory cytokines (*Ifng*, *Tnfα* and *Il1b*) analyzed by RT-qPCR, with Ct values normalized to *Actb*, in colon tissues of DSS- or water-treated mice fed on RCD, MFD, and RCD-MFD (n= 6). **K&L:** Expression levels of gut barrier genes (*Tjp1, Ocln, Cldn7,* and *Muc2*) (K) and AMPs (*Reg3b*, and *Lypd8*) (L) analyzed by RT-qPCR, with Ct values normalized to *Actb*, in IECs of mice, fed RCD, MFD, HFD, and RCD-MFD (n = 6). **M&N:** Western blot analysis of IECs isolated from mice fed on RCD, MFD, HFD, and RCD-MFD, showing p-JNK, JNK, p-p38, and p38 (n = 4). **O:** Microbial beta-diversity index, Chao index of fecal microbiome isolated from mice fed on RCD, RCD-MFD, and MFD (n = 5). **P:** Box plot showing reduced abundance of D. newyorkensis in the RCD-MFD group compared to the MFD group (n = 4). Data are mean ± SEM, with statistical analysis by two-way ANOVA with Tukey’s multiple comparison test (B, C, E, G, H and J), two-way ANOVA with Šídák’s multiple comparison test (F), two-way ANOVA with Uncorrected Fisher’s LSD (D), one-way ANOVA with Tukey’s multiple comparison test (K and L), and Mann Whitney test (P). *p < 0.05, **p < 0.01, ***p < 0.005, ****p < 0.001.

To further confirm this, we subjected one group of mice to a milk-based diet two weeks after weaning onto RCD (RCD-MFD), and another group to RCD two weeks after weaning onto MFD (MFD-RCD) (Figure S9D). Upon DSS treatment, both RCD-MFD and MFD-RCD mice exhibited reduced colon length and increased disease severity comparable to chow-fed mice, even though the loss of body weight was less drastic for the MFD-RCD group (Figure 5F–H & Figure S9E-G). Histopathology analysis of the RCD-MFD colon also showed increased pathology in these mice (Figure 5I). Pro-inflammatory cytokine mRNAs *Il1b*, *Ifng*, and *Tnfa* were also upregulated in the whole colon tissue of the RCD-MFD fed group, indicating increased inflammatory response (Figure 5J). Further, we specifically analysed the epithelial compartment without DSS challenge to see the basal expression of the tight junction proteins and anti-microbial peptides (*Reg3b* and *Lypd8*) and found them to be downregulated in the RCD-MFD group compared with the MFD group (Figure 5K and L). Interestingly, the RCD-MFD group showed expression of these genes similar to that in HFD-fed mice (Figure 5K and L). These results clearly show that unless the milk-based diets were initiated during the weaning phase, the signature changes associated with a protective milieu were abolished.

Further, we found that, unlike the MFD group, the RCD-MFD group did not show upregulation of JNK2 phosphorylation (Figure 5M), but they showed upregulation of P38 phosphorylation (Figure 5N). These data clearly indicate that a break in the milk-based diet feeding fails to activate JNK2 and alleviate its protective effect; instead, it activates the pro-inflammatory P38 pathway. We argued that this difference in the effect of MFD based on the time of initiation may reflect the context-dependent enrichment of distinct microbiomes by this diet. When initiated at 3 weeks, the peak of the weaning reaction, MFD enhances a microbiome characterised by high levels of *D. newyorkensis*, while late initiation does not favour the growth of this bacterium. Indeed, quantification of 16S rRNA sequencing data shows that the RCD-MFD microbiome differs from that of MFD in composition of species but not in species richness or evenness (Figure 5O and S9I). Additionally, RCDlllMFD exhibits lower levels of *D. newyorkensis* (Figures 5P and S9H), reduced concentrations of acetate and propionate in fecal matter, and no significant change in butyrate levels (Figure S9J). Taken together, our data show that continuation of milk supplementation immediately after weaning maintains a healthy microbiome, which enhances the gut barrier function by maintaining a microbiome enriched with *D. newyorkensis*, leading to activation of the JNK2 pathway in epithelial cells through acetate. On the other hand, if the milk-based diets are initiated at a later stage, the resultant microbiome is non-protective and activates the P38 pathway, facilitating inflammation.

## Discussion

Dietary modulation of the healthy microbiome is a promising strategy to address various conditions ranging from malnutrition to chronic disorders (*65*, *66*). It is established that diet is the primary determinant of the composition of the gut microbiome and its effect on gut pathologies (*10*). Nonetheless, our findings demonstrate that an identical diet can foster either a health-promoting or pathogenic microbiome, resulting in contrasting effects on gut pathology based on the context. Continued feeding of the pups with milk-based diets after weaning enhances gut barrier function upon DSS treatment, as evidenced by the maintenance of the gut epithelium in the milk-based diet-fed mice compared with RCD or HFD-fed mice. On the other hand, MFD, if initiated two weeks after weaning (RCD-MFD group), does not provide any protective effect. The deficiency of the latter condition is traced to its inability to induce a healthy microbiome characterised by high levels of *D. newyorkensis*. We argue that this may reflect the differences in the pre-existing microbiome at the time of MBD/MFD initiation. At the time of weaning, the gut microbiome is metastable (*4*), and the intervention with dairy establishes a microbial equilibrium that is protective. On the other hand, weaning to RCD establishes an alternative equilibrium with a different metabolic and inflammatory potential. Once the latter, non-protective microbiome is established, its response to MFD is clearly distinct from the response of the weaning stage microbiome, resulting in increased pathology. These results suggest that to determine the arc of the adult microbiome development, dietary modulation at the weaning stage is necessary. Extension of this study in humans would be useful in designing microbiome-directed approaches to reduce metabolic and inflammatory disease risks.

Because the diet-microbiome-inflammation relationship is indeed complex, the development of dietary intervention strategies requires a framework based on the associated molecular mechanisms. This study reveals a complex relationship between diet, gut microbiome, intestinal barrier, and inflammation shaped by post-weaning consumption of dairy. The key molecule that drives gut protection when MFD is initiated immediately after weaning is the short-chain fatty acid acetate, produced presumably by the microbiome, especially *D. newyorkensis*. Acetate, like other short-chain fatty acids, is known to enhance gut protection in similar models; however, the pathways by which it mediates protection were not well understood. Here, we show that activation of the JNK2 pathway in the colonic epithelium instructs enhanced gut barrier function, while activation of P38 is associated with increased inflammation.

JNK and P38 pathways are two core signalling pathways that control cell physiology, including cell proliferation, stress response, and inflammation (*12*, *14*). These pathways are well studied in the immune cells in the context of gut inflammation, and the JNK pathway has been shown to affect gut barrier function (*12*, *15*, *16*, *18*, *22*, *67*). However, how their functions are modulated by the microbiome to alleviate pathological gut inflammation was not known. Here, we suggest that these two pathways balance gut barrier integrity and inflammation. Milk-based diets, *D. newyorkensis*, and acetate activate JNK2 and reasonably maintain gut epithelial integrity even after a challenge with DSS for 7 days. On the other hand, feeding HFD, which induces low-grade inflammation even in the absence of any DSS challenge, leads to the activation of P38 in the epithelial cells. The RCD-MFD group also exhibited higher P38 and lower JNK2 phosphorylation. Moreover, inhibition of JNK activity upregulated P38 phosphorylation and alleviated the protective effects of milk-based diets, indicating that these pathways antagonise each other. Thus, we conclude that the relative levels of these pathways in the colonocytes define susceptibility to pathological inflammation in the gut. Our data shows that activation of the JNK2 pathway by acetate does not include the GPR43 activation. The possibility of epigenetic or metabolic remodelling of epithelial cells by acetate could be the underlying mechanism by which it activates JNK2, but it needs to be tested.

Two distinct features of gut protection downstream to the acetate-JNK2 axis are identified in the host: upregulation of gut barrier protection genes such as tight junction proteins, and the expression of antimicrobial proteins Reg3b and Lypd8. Tight junction proteins are essential in the maintenance of the gut epithelial barrier and are known to be upregulated by activation of transcription factors such as AP-1, HNF4α and CDX2, which are also upregulated by milk-based diets (*32*, *39*). Both REG3B and LYPD8 are the predominant AMPs expressed in the colon; on the other hand, a wider array of AMPs is expressed in the proximal intestine. REG3B and LYPD8 are gut protective as their deletion induces gut inflammation (*68*, *69*). Both REG3B and LYPD8 affect only a sub-population of microbes, including Gram-negative flagellated bacteria (*70*, *71*).

Interestingly, all these changes are affected by antibiotic treatment, suggesting that the protective effects are due to microbial products rather than a direct effect of dietary components on host cells. In support of this, all three features were restored when the mice were treated with *D. newyorkensis*. *D. newyorkensis* has been shown to enhance gut protection recently (*58*). The proposed mechanisms include immune modulation by maintaining Th17/Treg balance. Lysine produced by D. newyorkensis was shown to achieve immune tolerance by activating the kynurenine pathway in dendritic cells. Our data also showed that the protective myeloid populations were preserved in the MBD-fed mice treated with DSS, but emphasised the effect of this bacterium on the epithelial cells. The data presented advances the function of *D. newyorkensis* to enhance gut barrier function by activating the JNK2 pathway in the epithelium through acetate in mice.

The host controls the microbiome by secreting various molecules, including IgA, AMPs, and mucins (*11*). The microbiome controls the host and the microbial ecology by secreting various metabolites, antimicrobial peptides, etc. (*72*). The milk-based diets and *D. newyorkensis* treatment led to the upregulation of AMP genes *Reg3b* and *Lypd8*, which specifically target gram-negative and flagellated bacteria. However, antibiotic treatment abolished the effect of diet on the AMP expression, clearly suggesting the microbial dependence of their activation. Also, the lower expression of AMP coding genes in the RCD-MFD group, where the microbiome was different from the MFD group, suggests that the establishment of *D. newyorkensis* in the ecosystem as a predominant species would require induction of AMPs by the epithelial cells. The association of milk-based diets and *D. newyorkensis* treatment with AMP expression suggests that the abundance of *D. newyorkensis* could be determined by the feed-forward mechanism induced by this bacterium, mediated by AMPs, a process missing in the RCD-MFD group. This hypothesis needs to be tested in gene knock-out models of these AMPs.

While milk and dairy products are important nutritional ingredients, the effect of milk on gut health after weaning has been poorly understood. Milk and gut inflammation have a contentious relationship in adults. Milk and milk products remain one of the most commonly excluded dietary ingredients by patients with inflammatory bowel disease (*7–9*). However, clinical studies have not yielded an unequivocal conclusion on the adverse or protective effects of dairy (*8*, *9*, *73*). Some of the discrepancies in the clinical studies could be induced by the microbiome. Our data exposes the concept that the same diet could induce different outcomes based on the gut microbiome composition, as exemplified by the stark contrast between the effects of the MFD group and the RCD-MFD group on DSS-induced gut inflammation.

## Conclusion

In conclusion, our data indicate that the modulation of the microbiome from the weaning days by dietary modulation can have long-lasting effects on gut inflammation. More importantly, we also show that the effect of diets on the microbiome and inflammation depends on the pre-existing metabolic milieu and microbiome composition. These results are more relevant considering the association between malnutrition and dysbiosis and the effectiveness of early-life microbiome management in addressing growth defects associated with malnutrition (*2*, *5*). Further, the dichotomy of colonic epithelial JNK2 and P38 activation predetermines the response of gut epithelium to insults that weaken gut barrier function. While activation of JNK2 is essential for enhanced gut epithelial integrity by acetate, P38 activation is associated with increased inflammation.

## Materials and Methods

### Materials

Chemicals, reagents, kits, strains and software used in this study are given in tables S3-S8.

### Animal maintenance

C57BL/6 mice (Jackson Laboratory, cat. no. 000664) were obtained and bred in the small animal facility at the National Institute of Immunology, New Delhi, India. Male littermate mice were used for experiments and fed ad libitum with one of the following diets: a standard rodent chow diet (RCD; Altromin, cat. no. 1324), a Lactogen-2 paste (Nestlé) (MBD), a formulated milk fat diet (MFD, Research Diets, cat. no. D19112203), or a high fat/Western diet (HFD; Research Diets, cat. no. D12492I). All animals were housed in individually ventilated cages (IVCs). Pups were weaned onto one of the designated diets at 24 days of age. Each experimental group included a minimum of three mice, and each experiment was repeated at least twice. All procedures involving animal care, strain maintenance, dietary interventions, post-mortem tissue collection, and drug treatments were conducted in accordance with institutional guidelines and were approved by the Animal Ethics Committee of the National Institute of Immunology.

### Dextran sodium sulphate-induced colitis

Mice were weaned onto four different dietary regimens: (1) RCD, (2) MBD, (3) MFD, and (4) HFD as described above. After 28 days of feeding (at 8 weeks of age), we further divided each group into two subgroups: (a) treated mice that received either 1.5% or 2.5% DSS (MFD and RCD groups only) in drinking water, and (b) control mice that received water without DSS. For the 1.5% DSS group, each experimental group consisted of 4 animals each, and the experiment was independently repeated three times (n = 12 per group). For the 2.5% DSS group, the control group included 5 animals (n = 5), and the DSS-treated group included 7 animals (n = 7). We administered DSS for 7 days and monitored the body weight and disease severity daily. We scored disease severity based on stool consistency and clinical symptoms: 0 points for well-formed pellets, 1 point for pasty or semi-formed stool, 2 points for liquid stool, 3 points for stool with a bloody smear, and 4 points for bloody fluid or mortality (*74*). We classified mice that lost more than 30% of their body weight as moribund, euthanised, and excluded from the analysis. After euthanasia, we recorded colon length and collected colon tissues for downstream analyses. In the RCD-MFD group (n = 6), we fed mice RCD for 2 weeks post-weaning, then switched them to MFD for 2 weeks. Meanwhile, in the MFD-RCD group (n = 3), we fed mice MFD for 2 weeks post-weaning, then switched them to RCD for two weeks. We allowed a 2-week recovery period between two 7-day DSS cycles for repeated exposure.

### 2,4,6-trinitrobenzenesulfonic acid (TNBS) induced colitis

Mice were weaned onto two dietary regimens: (1) RCD, and (2) MFD. After 28 days of feeding (at 8 weeks of age), we further divided each group into two subgroups: (a) treated mice that received 2.5% TNBS, and (b) control mice. The control group included 5 animals (n = 5), and the 2.5% TNBS-treated group included 8 animals (n = 8). We applied 150 µL of 1% (wt/vol) TNBS to the shaved dorsal skin of 7-week-old mice for presensitization. Eight days later, we recorded the body weight and fasted the mice for 12 hours with free access to water. For colitis induction, we anaesthetised the mice with an intraperitoneal injection of ketamine (100 mg/kg) and xylazine (10 mg/kg), then administered 100 µL of 2.5% TNBS in 50% ethanol intrarectally using a catheter inserted 3.5 cm into the rectum as described (*75*). We held the mice in a Trendelenburg position for 5 minutes after instillation. We monitored clinical signs of colitis daily, including changes in body weight, stool consistency, and rectal bleeding. We classified mice that lost more than 30% of their body weight as moribund, euthanised, and excluded from the analysis. We euthanised the mice on 5^th^ day of treatment and collected samples for downstream analyses.

### FITC-Dextran cell permeability assay

Mice were weaned onto three different dietary regimens: (1) RCD, (2) MBD, and (3) MFD, and subjected to DSS-induced colitis as described above. Each experimental group consisted of 3 animals, and the experiment was independently repeated two times (n = 6). On day 7 of treatment, we fasted the mice for 2 hours and then administered intra-gastric FITC-dextran (0.5 mg/g body weight). Six hours later, we collected blood for serum analysis to assess intestinal permeability. We measured the FITC fluorescence in the serum using a SpectraMax M2 plate reader, with excitation at 490 nm and emission detected at 510 nm.

### Oral administration of Acetate

We supplemented the drinking water of 4-week-old mice on RCD with 200 mM potassium acetate for three weeks, followed by treatment with 1.5 % DSS containing acetate (n = 8).

### JNK inhibition in the colon

We anaesthetised 8-week-old RCD and MFD fed mice (n = 8) with an intraperitoneal injection of ketamine (100 mg/kg) and xylazine (10 mg/kg) at a volume of 40 µL per mouse, following a 24-hour fasting period to clear the colon. We carefully inserted a catheter into the colon until the tip reached 4 cm proximal to the anus. We then administered 100 µL of JNK inhibitor II (10 µg/g; SP600125, Merck) in PBS containing 10% FBS. Control mice received the vehicle alone (PBS with 10% FBS). Following administration, we kept the mice in a Trendelenburg position for five minutes. We then allowed the mice to eat ad libitum and isolated the intestinal epithelial cells (IECs) at 6 and 24 hours post-treatment to confirm pathway inhibition. We initiated 1.5% DSS treatment 24 hours post-inhibitor administration.

### GPR43a inhibition in the colon through the rectal route

To inhibit GPR43a in the colon, we anaesthetised 8-week-old RCD and MFD fed mice (n = 4) with an intraperitoneal injection of ketamine (100 mg/kg) and xylazine (10 mg/kg) at a volume of 40 µL per mouse, following a 12-hour fasting period. We inserted a catheter 4 cm into the colon via the anus and administered 100 µL of GPR43a inhibitor (10 µg/g; GLPG0974, Merck) in PBS containing 0.4% DMSO. Control mice received the vehicle alone (PBS with 0.4% DMSO). Following administration, we placed the mice in a Trendelenburg position for 5 minutes. To ensure effective inhibition, a second dose was administered after 2 days. We isolated the intestinal epithelial cells (IECs) for downstream analysis six hours after the final dose.

### Antibiotic treatment

To deplete gut microbiota, RCD, MFD and HFD fed mice (n = 6) were given a broad-spectrum antibiotic cocktail in drinking water from 5 to 7 weeks of age. The cocktail contained Vancomycin (0.5 mg/ml), Ampicillin (1 mg/ml), Neomycin (0.5 mg/ml), and Metronidazole (0.625 mg/ml), in 2% sucrose. Continuous access to antibiotic-containing water was maintained throughout the treatment. To confirm microbial depletion, 50 mg of fecal matter from ABX-treated and control mice was resuspended in 1 ml PBS, serially diluted (1/4 and 1/8), and plated for microbial growth assessment.

### Faecal matter transplant

We performed faecal microbiota transplantation (FMT) by collecting fresh faecal pellets from healthy donor RCD and MFD fed mice (n = 8) housed under specific pathogen-free conditions. We homogenised the faeces in sterile 1× PBS, followed by centrifugation of the suspension at 800 × g for 3 minutes to remove large debris. We then filtered the supernatant through a 40 µm cell strainer. Before FMT, we treated the recipient RCD and MFD fed mice (n =8) with 200 µL of PEG 6000 at a concentration of 425 g/L, as described by (*76*) to enable microbiota clearance. Four hours after microbiota clearance, we administered 200 µL of the filtered faecal suspension containing 20 mg of faecal material per 1 mL of PBS via oral gavage to each recipient mouse every two days for three weeks. Mice in the control group received 200 µL of sterile 1× PBS following the same schedule. We monitored all mice for general health and treated them with 1.5% DSS.

### Real-time qPCR analysis of bacterial 16S rRNA genes

We collected feces from antibiotic-treated mice and from each mouse (n = 6) 6 hours before and after PEG treatment, then immediately stored the samples at −80LJ°C. We extracted total DNA from 100 mg of feces using the QIAamp® Fast DNA Stool Kit (Cat. No. 51604) and stored the DNA at −20LJ°C. We performed real-time qPCR targeting 16S ribosomal genes using SYBR® Premix Ex Taq PCR master mix on a QuantStudio™ 6 Flex Real-Time PCR System. We compared the cycle threshold (Ct) values of each sample within their respective groups and represented them as an amplification plot.

### Administration of *Duboisiella newyorkensis* as a probiotic

*Duboisiella newyorkensis* was obtained from the American Type Culture Collection (ATCC, No. TSD-64-0.5ML) and cultured in modified Tryptone Glucose Meat extract (MTGE) broth at 37 °C under anaerobic conditions. We estimated the concentration of the bacterial culture by measuring the optical density at 600 nm (OD600). We used bacterial cultures with an OD of 0.2-0.4 for treatment. We treated 5-week-old RCD fed mice (n = 8) with a broad-spectrum antibiotic cocktail to deplete the gut microbiota for one week. 100µl of *D. newyorkensis* (1OD/ 100 µl) culture was administered via oral gavage once every two days for three weeks, followed by treatment with 1.5% DSS. Control mice received 100 µL of sterile MTGE media following the same schedule.

### Histological analysis using Haematoxylin and Eosin staining

We excised the distal colon tissues from mice after euthanasia and washed them with ice-cold phosphate-buffered saline (PBS, pH 7.2). We embedded the tissues in tissue-freezing media (Cat. SHH00260) and snap-frosted them in liquid nitrogen. Using a cryotome, we prepared cryosections at a thickness of 10-μm. We then stained the sections with hematoxylin and eosin and captured images using an Olympus inverted microscope with a 20X objective, with image acquisition performed using Image-Pro 6 software.

### Intestinal epithelial cell (IEC) isolation

We euthanised the mice, carefully dissected the colon, and cut it open longitudinally. We washed the tissue three times with PBS, cut it into small pieces, and incubated it in a pre-digestion buffer (1X PBS with 10 mM EDTA, 0.5 mM DTT) at 37°C with shaking at 200 RPM for 15 minutes. After incubation, we removed the pre-digestion buffer by centrifugation and added a wash medium (RPMI + 5% FBS). We vortexed the tissue suspension to release epithelial cells, collected the supernatant and filtered it through a nylon cell strainer with a 70-μm pore size.

### RNA Isolation and RT-PCR

We isolated RNA from approximately 0.5 cm of distal colon tissue or IECs using the Macherey-Nagel Kit (Cat. no. 740955.50). We removed genomic DNA by treating the samples with DNase provided in the kit. We quantified the purified RNA using a spectrophotometer (Multiscan SkyHigh) and synthesised cDNA using the Takara cDNA synthesis kit (Cat. no. 6110A). We performed qRT-PCR using SYBR® Premix Ex Taq PCR master mix (Takara Cat. # RR420A) on a QuantStudio™ 6 Flex Real-Time PCR System. We calculated relative mRNA expression levels using the 2^−ΔΔCt^ method, using actin as the internal control.

### Western Blot Analysis

We lysed isolated IECs or colon tissues using RIPA lysis buffer and a homogeniser, then centrifuged the lysates. We measured the protein concentration in the supernatants using a BCA assay (Thermo Scientific, catalogue no. 23227). We resolved 40 μg of total protein by SDS-PAGE and transferred the proteins onto a nitrocellulose membrane (Cytiva, Catalogue no. 10600002). We probed the membrane with primary antibodies against p38, phospho-p38, SAP/JNK, phospho-SAP/JNK, cyclinD1, occludin, MKK4, MKK7, cleaved caspase3, and β-actin. We used corresponding HRP-conjugated secondary antibodies and developed the bands using ECL reagent. We captured the images using the Azure biosystems 300 chemidoc system.

### Ki67 staining of colon tissue cryosections

We equilibrated frozen cryosections to room temperature (RT) for 30 minutes, fixed them in 4% PFA for 20 minutes, and performed antigen retrieval at 100°C for 20 minutes using sodium citrate buffer. After cooling and washing, we blocked the tissues with 200 μL blocking buffer (1 X PBST + 2% BSA + 5%BSA) for 1 hour at RT and incubated them overnight at 4°C with Ki67 primary antibody (1:400). We then washed the slides and incubated them with secondary antibody for 1 hour at RT, followed by three washes. After air drying, we added 10 μL Fluoroshield and carefully mounted coverslips and captured images using an ZEISS Axio Imager 2 microscope with a 20X objective, with image acquisition performed using AxioVison (V 4.8.2.0)

### Lamina propria immune cell isolation

We euthanised the mice, carefully dissected the colon, and cut it open longitudinally. We washed the tissue three times with PBS, cut it into small pieces, and incubated it in a pre-digestion buffer (1X PBS with 25 mM HEPES, 10 mM EDTA, 0.5 mM DTT) at 37°C with shaking at 200 RPM for 15 minutes. Following incubation, we centrifuged the mixture to remove the pre-digestion buffer and added wash medium (RPMI with 5% FBS). We vortexed the sample and collected the epithelial cells from the supernatant. We filtered the supernatant sequentially through nylon cell strainers with 70-μm and 40-μm pore sizes. We subsequently treated the remaining tissue with the digestion buffer (RPMI supplemented with 10% FBS, 25 mM HEPES, 0.5 mg/ml Collagenase IV, 0.5 mg/ml Dispase) with shaking at 200 RPM for 45 minutes at 37°C. After digestion, we centrifuged the mixture at 1500 RPM for 5 minutes at room temperature and discarded the supernatant. We added the pre-warmed wash medium, vortexed the sample, and filtered the supernatant sequentially through 70-μm and 40-μm nylon cell strainers. Finally, we counted the isolated cells.

### Cell sorting for single-cell RNAseq

We isolated epithelial and immune cells and determined the number of live cells using the Luna-II Automated Cell Counter. We pooled cells from three mice per group: RCD with water, RCD with 1.5% DSS, MBD with water, and MBD with 1.5% DSS in equal proportions. Using a BD FACSAria cell sorter, we sorted the cells into four populations: epithelial cells (EpCAM+), B cells (CD45+ CD19+), T cells (CD45+ CD3+), and myeloid cells (CD45+ CD3− CD19−). We pooled 6000 epithelial cells, 2000 B cells, 4000 T cells, and 8000 myeloid cells for single-cell RNA sequencing using the 10X Genomics platform for each group.

### 10X Genomics scRNAseq library preparation

We partitioned single cells into Gel Beads in Emulsion (GEMs) using the Chromium Controller and the Single Cell 3′ v3.1 Gel Beads Kit (10X Genomics, Cat. No. 2000164), following the manufacturer’s protocol. We performed GEM reverse transcription using the following PCR conditions: 53 °C for 45 minutes, 85 °C for 5 minutes, and a hold at 4 °C. Reverse transcription cleanup was carried out using Dynabeads MyOne SILANE. We amplified the resulting cDNA using PCR with the following conditions: 98 °C for 3 minutes, 11 cycles of 98 °C for 15 seconds, 63 °C for 20 seconds, 72 °C for 1 minute, followed by a final extension at 72 °C for 1 minute. We purified the amplified cDNA using Ampure XP beads and quantified it using a Fragment Analyser. We then performed fragmentation, end repair, and A-tailing, followed by double-sided size selection using Ampure XP beads. To construct the 3′ gene expression (GEX) library, followed by double-sided size selection using Ampure XP reagent. We ligated the GEX adaptors and cleaned them with Ampure XP beads. We subsequently performed GEX sample index PCR and carried out an additional round of double-sided size selection with Ampure XP. We assessed the quality of the final library using the Fragment Analyser before sequencing.

### Single-cell RNAseq analysis

Cell Ranger (v7.1.0) aligned the FASTQ files to the mouse reference genome and generated feature-barcode matrices. We performed downstream analysis using the Seurat package (v4.0). We retained cells that met the following quality control criteria: nCount_RNA > 1000, nFeature_RNA > 500, and singlets identified using DoubletFinder. A total of 38,922 cells passed QC and were included in the analysis. We normalised the data using SCTransform, setting var.to.regress to MitoPercent. We then performed Dimensionality reduction and clustering using the Seurat functions RunPCA, FindNeighbours, FindClusters, and RunUMAP. We used Seurat’s plotting function, the scCustomize package and in-house custom scripts for data visualisation. As described in the Results section, we manually annotated cell types based on canonical marker gene expression. We then subset individual cell types and reclustered them following the same pipeline. For regulome analysis, we applied the SCENIC package. The code used for this analysis will be made available upon request.

### Cecal DNA isolation and 16S rRNA gene sequencing on the MinION™ platform

We euthanised the mice and isolated the cecal content from RCD, MBD, MFD, HFD and RCD-MFD fed groups. We pooled 200 mg of cecal matter from three mice as one biological sample, with five samples per group. We extracted the cecal matter DNA using a combinatorial method consisting of chemical and mechanical lysis, as described by Shuming group (77). Following the manufacturer’s protocol, we prepared the full-length 16S rRNA library using the 16S barcoding kit 1-24 (SQK-16S024, Oxford-Nanopore Technologies). The amplified products were purified using AMPure® XP beads and quantified using Qubit™ dsDNA HS and BR Assay Kits (Thermo Scientific, Catalogue number: Q32854). 100 ng of DNA was incubated with 1 μl of Rapid Adapter at room temperature for five minutes. 11 μl of library was mixed with 34 μl of Sequencing Buffer, 25.5 μl of Loading Beads, and 4.5 μl of water, loaded onto the FLO-MIN106 (Oxford Nanopore Technologies), and sequenced on the MinION™. MINKNOW software ver. 1.11.5 (Oxford Nanopore Technologies) was used for data acquisition. Guppy basecaller (Oxford Nanopore Technologies) was used for base calling the MinION™ sequencing data (FAST5 files) to generate pass reads (FASTQ format) with a mean quality score > 7. The fastq files were aligned to the NCBI 16S rRNA dataset using EPI2ME tools. We generated abundances based on barcode demultiplexing and filtered them using the following quality control criteria: read length > 1000 bp and accuracy > 95%. Using the microeco R package, we calculated relative abundance and alpha and beta diversity metrics.

### Quantification of faecal SCFA using GC-MS

For metabolite extraction, 50 mg of caecal matter was mixed with 1μL of ribitol (10 mg/ml), as an internal standard and 500 μL of GC-MS grade water, followed by centrifugation. 10 µl of concentrated sulfuric acid (H_2_SO_4_) was combined with the 100 µl of extract, and 400 µl of diethyl ether (DE) was added to form a liquid-liquid extraction. 100 µl of DE extract is added to a GC vial. 1µl of BSTFA is added and vortexed for 5 seconds, followed by derivatisation at 37 °C for 2 hours (*78*). 1ul of derivatised samples was loaded to GC/MS (Agilent 7890A) coupled with 5975C MSD with Triple-Axis Detector (TAD). The injector, ion source, quad helium carrier gas flow rate, and temperatures were set to 260, 230, 150, and 280 °C, respectively. The helium carrier gas flow rate was kept at 1 ml/min. 1μl of the derivatized sample was injected with a 3 min of solvent delay time and split ratio of 10:1. The initial column temperature was 40 °C and held for 2 min, ramped to 150 °C at the rate of 15 °C/min and held for 1 min, and then finally increased to 300 °C at the rate of 30°C/min and kept at this temperature for 5 min. The ionisation was performed in the electron impact (EI) mode at 70 eV. The MS data were acquired in full scan mode from m/z 30–400 at an acquisition frequency of 12.8 scans per second. Comparison with pure standards, based on both retention time and the corresponding MS spectra, confirmed the identification of compounds.

### Statistics and reproducibility

All data are represented in a format that shows the data distribution clearly (dot plots) and all the graph elements (median and error bars) are defined in the figure legends. Sample size was not pre-determined by any statistical tests. In all figures and figure legends, *n* represents the number of experimental units. Statistical comparisons were performed using an ANOVA test with Dunnett’s, Tukey’s or Sidak’s correction for multiple samples, or using an unpaired Student’s *t*-test for two samples. The exact *P* values are indicated in the relevant graphs. The number of biological replicates is indicated in Figures.

## Ethical Standards Statement

The authors of this manuscript certify that all experiments in the study were approved by the Institutional Animal Ethics Committee.

## Acknowledgments

We are grateful to Mr. B. N. Roy for his help with histopathology experiments and Mr. Khem Singh Negi for technical help. We thank Dr. Nagarajan for his help with animal experiments. We thank Dr. Ranjan Nanda for providing Ribitol and discussions on GC-MS analysis. We thank Mr. Somdeb Chattopadhyay for running the SCENIC script and providing the results.

---

**Figure S1:**
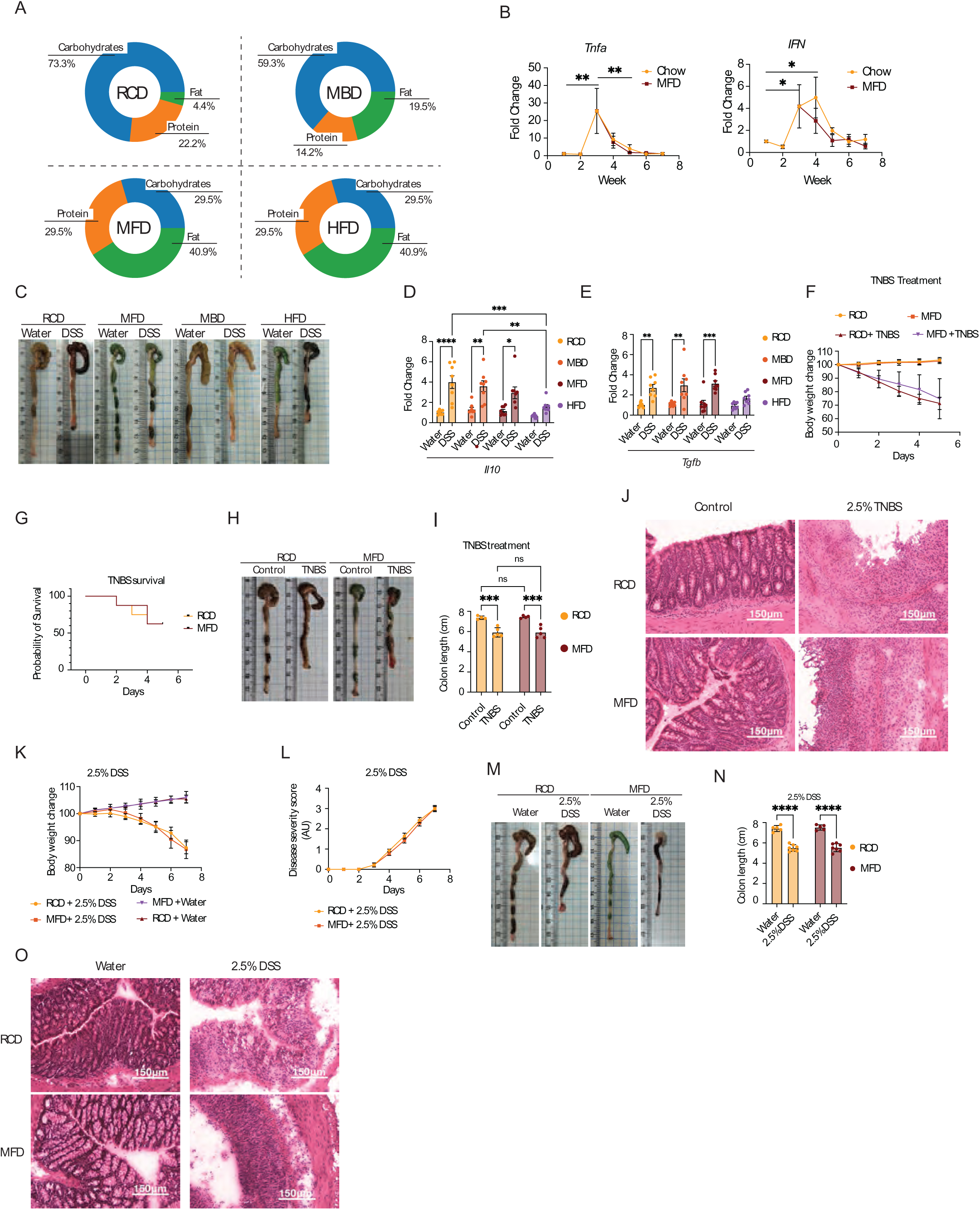
Higher dose of DSS and TNBS diminish the protective effects of milk-based diets. **A:** Composition of macronutrients in different diets: RCD, MBD, MFD, and HFD. **B:** RT-qPCR analysis of Tnfα and Ifnγ expression in colon tissues from mice fed RCD or MFD at different time points. Ct values were normalized to *Actb*. **C:** Representative images of the colon from DSS or water-treated mice fed RCD, MBD, MFD, and HFD. **D&E:** Expression levels of anti-inflammatory cytokines *Il10* (D) and *Tgfβ* (E) analyzed by RT-qPCR, with Ct values normalized to *Actb* mRNA, in colon tissues of DSS or water treated mice fed on RCD, MBD, MFD, and HFD (n = 8). **F-I:** Body weight changes (n = 5) (F), survival probability (n = 8) (G), representative images of the colon tissues (H), and total colon length in cm (n = 5) (I) mice treated with 2.5% TNBS and control either fed on RCD or MFD diets. **J:** Representative images of H&E stained colon tissues from TNBS treated or control mice either fed on a RCD or a MFD diets. (Scale bars: 100 μm). **K-N:** Body weight changes (K), disease severity score (L), representative images of the colon tissues (M), and total colon length in cm (N) of mice treated with 2.5% DSS either fed on a RCD or a MFD diets (n = 7). **O:** Representative images of H&E stained colon tissues from 2.5% DSS treated or control mice either fed on a RCD or a MFD diets. (Scale bars: 150 μm). Data are mean ± SEM of two independent experiments, with statistical analysis by two-way ANOVA with Tukey’s multiple comparison test (B, D, E, F, I, K, and L), Mixed-effects analysis with Tukey’s multiple comparison test (B *Ifnγ*), two-way ANOVA with Šídák’s multiple comparison test (N), and Survival test (G). *p < 0.05, **p < 0.01, ***p < 0.005, ****p < 0.001.

**Figure S2:**
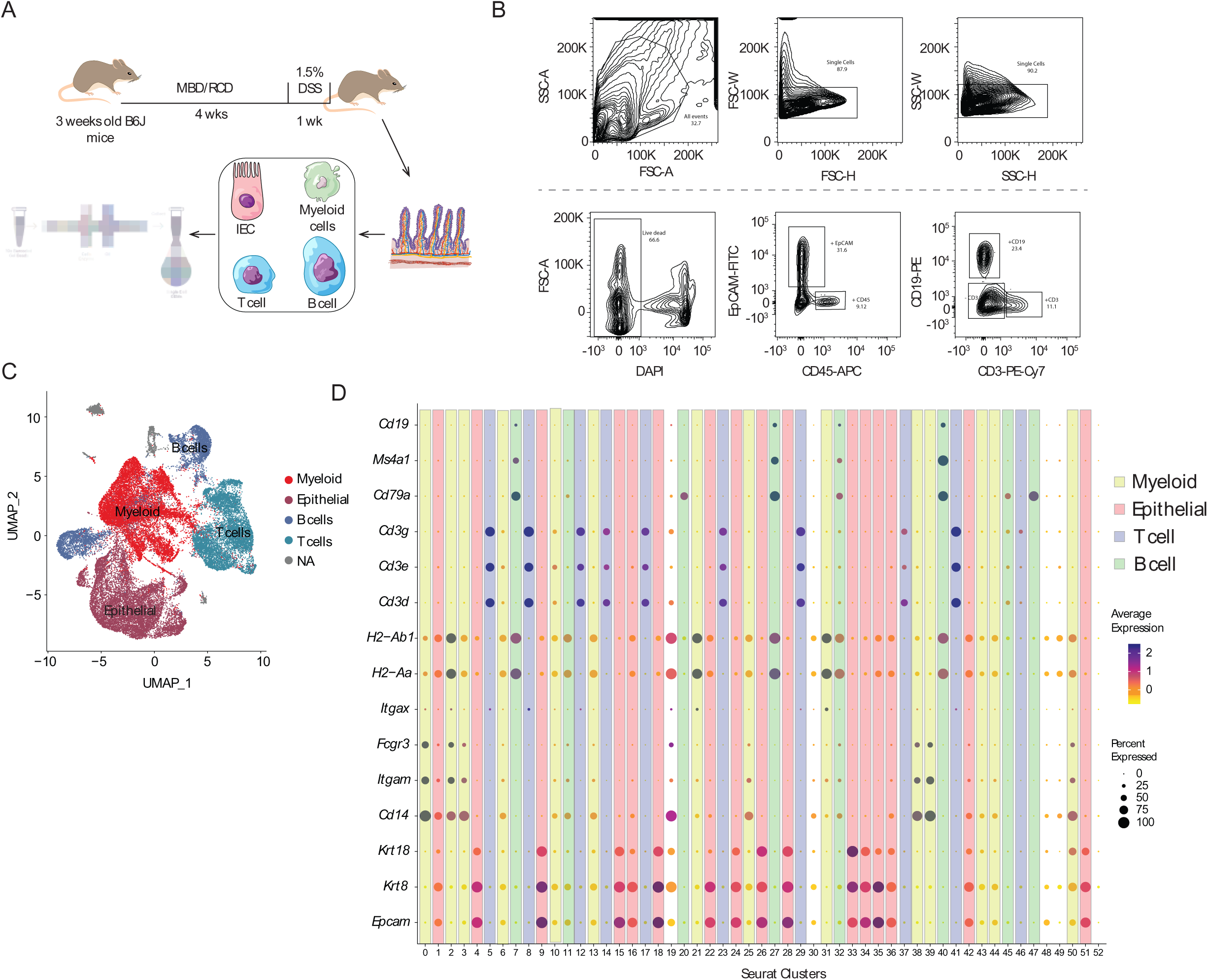
scRNA-seq gating strategy and expression of cell type–specific markers. **A:** Schematic of the scRNA-seq workflow. Mice were weaned onto MBD or RCD for 4 weeks, then treated with water or 1.5% DSS for 7 days. Colons were harvested, and IECs, myeloid cells, T cells, and B cells were isolated, pooled, and processed for scRNA-seq. **B:** Gating strategy for sorting epithelial, T cell, B cell, and myeloid cells from colon. **C:** UMAP representation of all detected cells, highlighting epithelial cells, B cells, T cells, and myeloid cells based on markers. **D:** Dot plot of marker genes highlighting epithelial cells (*Epcam, Krt8,* and *Krt18*), B cells (*CD79a, Ms4a1,* and *CD19*), T cells (*CD3d, CD3e,* and *CD3g*), and myeloid cells (*CD14, Itgam, Fcgr3, Itgax, H2-Aa,* and *H2-Ab1*). The color indicates the expression level, and the size of the dots indicates the percentage of cells expressing the gene.

**Figure S3:-.**
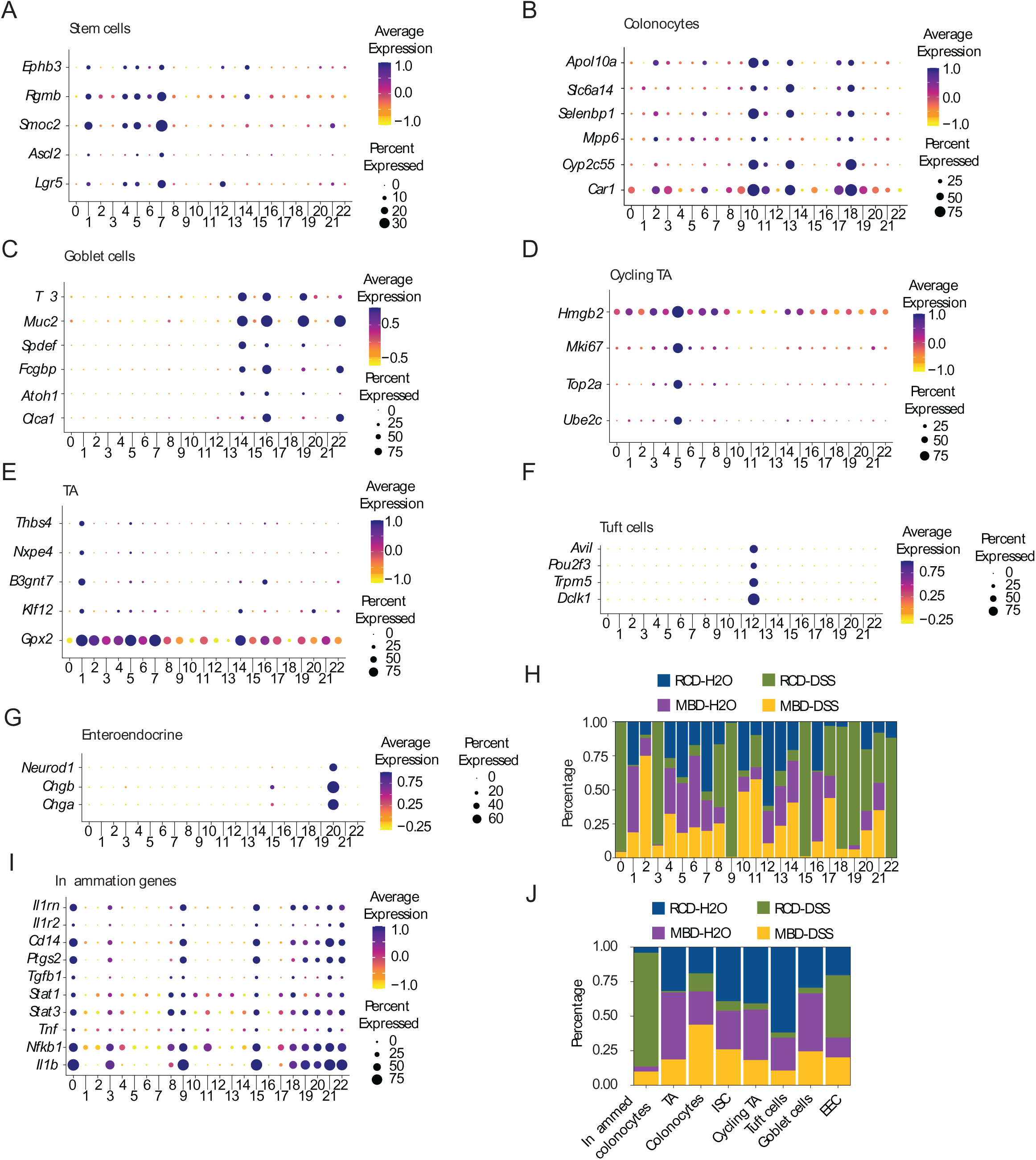
Cell type–specific marker expression and cluster distribution across experimental groups in epithelial cells. **A-G:** Dot plots showing expression levels of marker genes for stem cells (A), colonocytes (B), goblet cells (C), cycling TA (D), TA (E), tuft cells (F), and enteroendocrine cells (G) across all epithelial clusters, colored by average gene expression. Dot size indicates the percentage of cells expressing the gene. **H&J:** The percentage of cells from each experimental group falling into different clusters (H) or cell types (J) were plotted as a stacked bar plot. Each bar is standardized to have the same height using the position_fill() argument in ggplot2. Salmon represents RCD-H2O, green represents RCD-DSS, dark turquoise represents MBD-H2O, and violet represents MBD-DSS. **I:** Dot plots showing expression levels of inflammatory genes across epithelial cell clusters. Color intensity indicates the average expression level, and the size of the dot indicates the percentage of cells expressing the gene.

**Figure S4:-.**
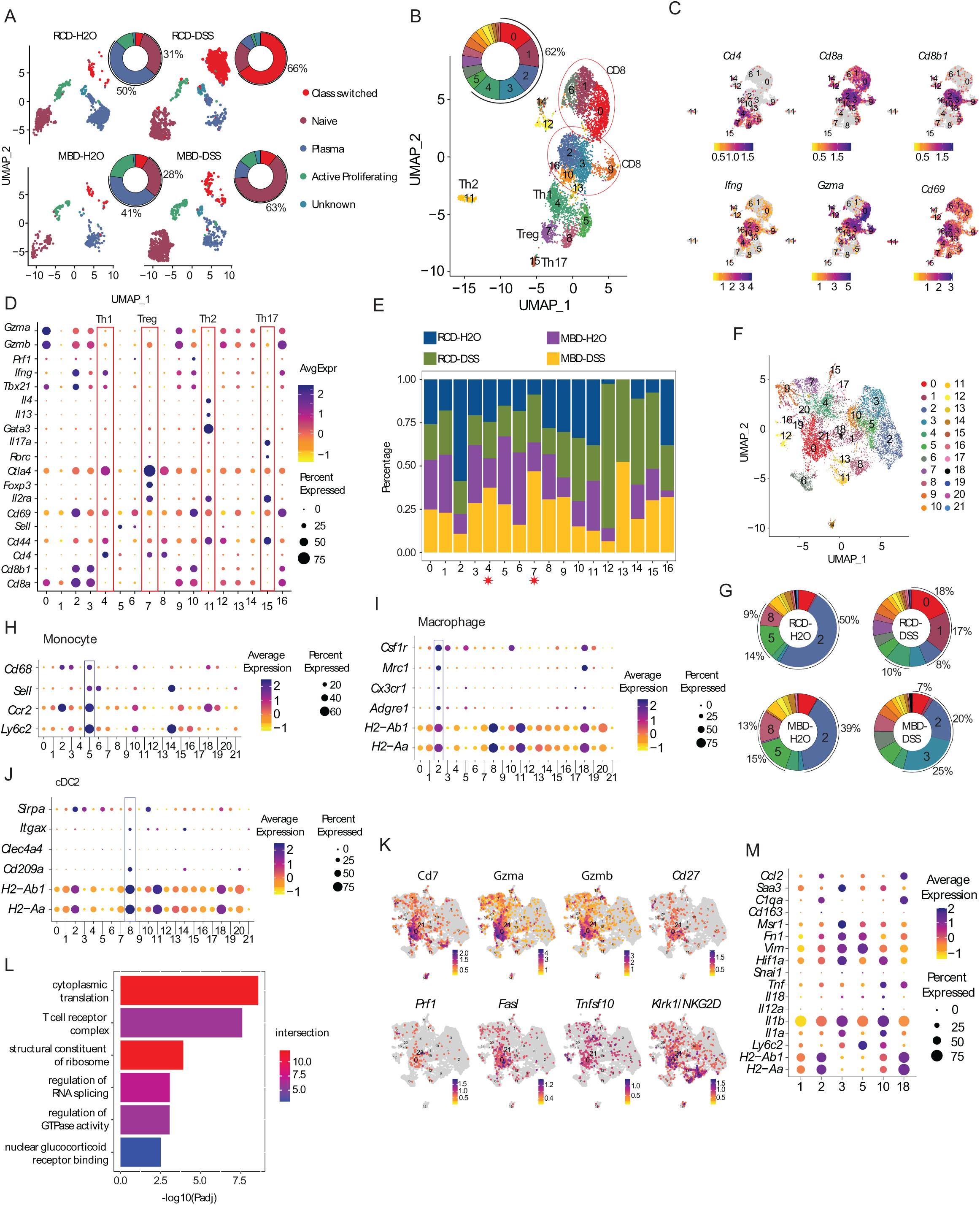
Single-cell transcriptomic profiling of B cells, T cells, and myeloid cells across dietary and DSS treatment conditions. **A:** UMAP and donut plots showing B cell distribution across cell types in RCD-water, RCD-DSS, MBD-water, and MBD-DSS groups. Percentages for selected cell types are indicated. **B:** UMAP and donut plot showing T cell distribution across clusters with selected cell type percentages indicated. **C:** UMAP plots showing expression of T cell markers (*Cd4, Cd8a, Cd8b1, Ifng, Gzma, Cd69)*, with color intensity indicating expression levels. **D:** Dot plot showing expression of various T cell markers, highlighting different subsets such as Treg, Th1, Th2, and Th17 cells. The color indicates the expression level and the size of the dots indicates the percentage of cells expressing the gene. **E:** The percentage of cells from each experimental group falling into different clusters was plotted as a stacked bar plot. Each bar is standardized to have the same height by using the position_fill() argument in ggplot2. Dark blue represents RCD-H2O, green represents RCD-DSS, violet represents MBD-H2O, and orange represents MBD-DSS. **F:** UMAP representation of all myeloid cells (*Cd14, Itgam, Fcgr3, Itgax*) showing 22 clusters. **G:** Donut plots show the distribution of myeloid cells from different experimental groups across different clusters. The percentage of cells belonging to some of the clusters is shown next to each donut plot. **H-J:** Dot plots showing expression levels of monocyte (H), macrophage (I), and cDC2 (J) marker genes. The color indicates the expression level, and the size of the dots indicates the percentage of cells expressing the gene. **K:** UMAP plots colored by the expression of cytotoxic NK cell marker genes (*Klrk1/NKG2D, Cd7, Cd27, Gzma, Gzmb, Prf1, Fasl,* and *Tnfsf10*). The intensity of the color indicates the expression levels as shown in the color scale given with each plot. **L:** Bar plot showing −log10(adjusted p-value) of GO terms enriched in cluster 0 of myeloid cells. The color indicates the number of genes differentially expressed in cluster 0 that overlaps with the indicated GO term. **M:** Dot plot showing expression of various macrophage and inflammatory monocyte marker genes. The color indicates the expression level, and the size of the dots indicates the percentage of cells expressing the gene.

**Figure S5:-.**
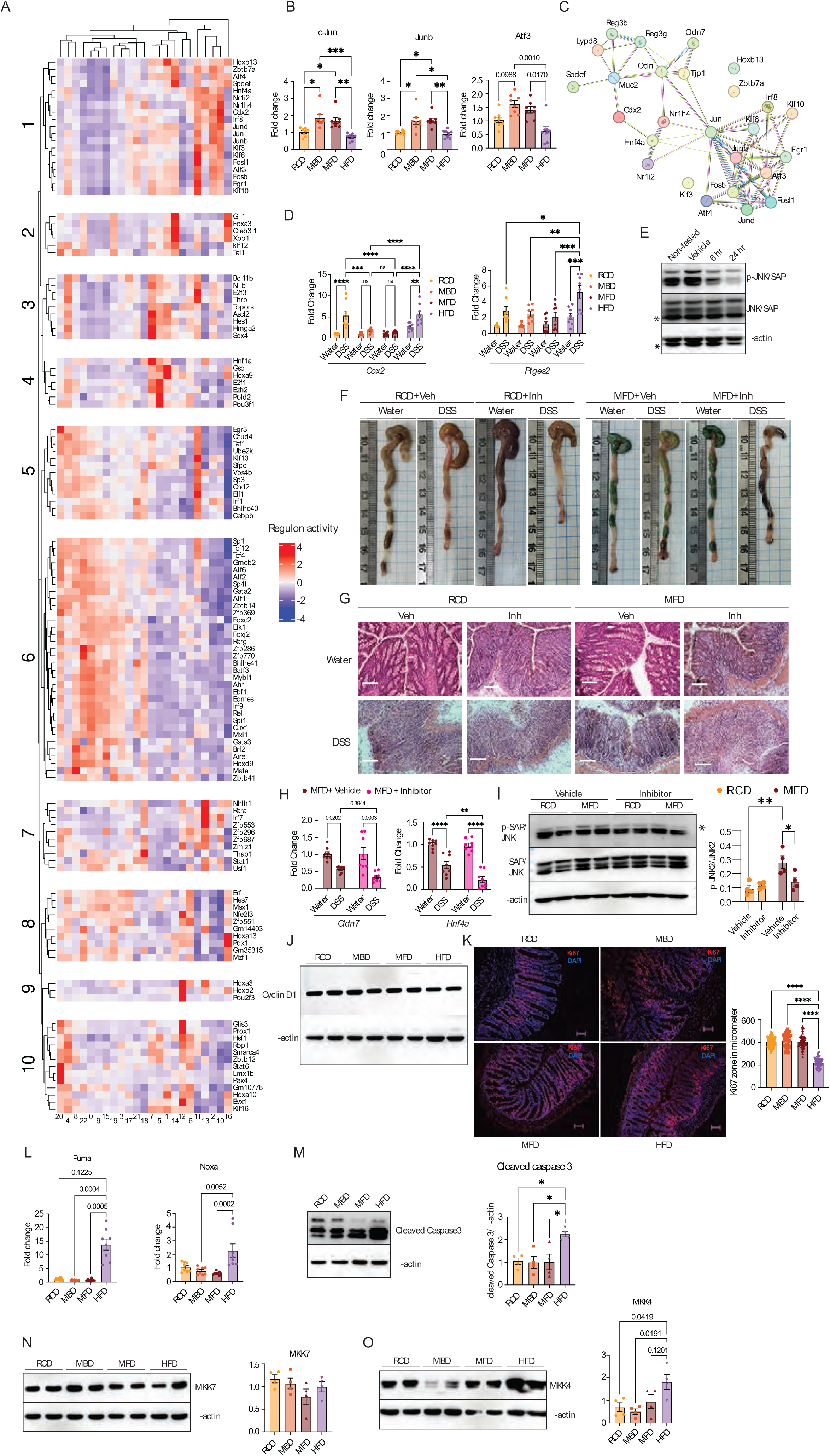
JNK-dependent responses to diet and DSS in the colon tissue. **A:** Heatmap showing the regulome expression calculated using the SCENIC package. **B:** JNK downstream genes associated *cJun, Junb* and *Atf3* analyzed by RT-qPCR, with Ct values normalized to *Actb*, in colon tissues of DSS- or water-treated mice, fed RCD, MBD, MFD, and HFD (n = 8). **C:** Network graph showing the interaction between transcription factors identified in the SCENIC analysis and gut barrier and AMP genes analyzed by StringDB. **D:** p-p38 target genes associated with inflammation (*Cox2*, and Ptges2) analyzed by RT-qPCR, with Ct values normalized to *Actb*, in colon tissues of DSS- or water-treated mice, fed RCD, MBD, MFD, and HFD (n = 8). **E:** Western blot analysis for p-JNK and JNK levels in IECs isolated from JNK inhibitor II treated mice fed on RCD. Samples were collected at 6 hrs and 24 hrs post inhibitor treatment. **F:** Representative images of colon tissues, DSS or water-treated, with or without JNK inhibitor II. Diet groups are indicated. **G:** Representative images of H&E-stained colon tissue sections of RCD and MFD-fed mice treated with DSS- or water and subjected to JNK inhibition or vehicle treatment. (Scale bars: 100 μm). **H:** Gut-barrier associated genes *Cldn7* and *Hnf4α* analyzed by RT-qPCR, with Ct values normalized to *Actb*, in colon tissues of DSS- or water-treated mice with or without JNK inhibition, fed on MFD (n = 8). **I:** p-JNK and JNK levels were analyzed in IECs from mice fed on RCD and MFD, isolated 7 days after p-JNK inhibition (n = 4). Representative blot (left panel) and densitometric quantification (right panel) are shown. **J:** Western blot analysis of Cyclin D levels in IECs from mice fed RCD, MBD, MFD, or HFD (n = 4). **K:** Ki67 staining of colon cryosections showing the proliferation zone (left panel). Quantification of proliferation zone length in µm (right panel). **L:** RT-qPCR analysis of apoptosis-associated genes *Puma* and *Noxa* in colonic IECs from mice fed RCD, MBD, MFD, or HFD (n = 8). Ct values were normalized to *Actb*. **M-0:** Western blot analysis of cleaved-caspase3 (M), MKK7 (N), and MKK4 (O) levels in IECs from mice fed RCD, MBD, MFD, or HFD (n = 4), with quantification shown in the right panel. Data are mean ± SEM, with statistical analysis by Kruskal-wallis test with Dunn’s multiple comparisons test (B, and L), two-way ANOVA with Tukey’s multiple comparison test (D, H, and I) and one-way ANOVA with Holm-Šídák’s multiple comparison test (K, M, N, and O). *p < 0.05, **p < 0.01, ***p < 0.005, ****p < 0.001.

**Figure S6:**
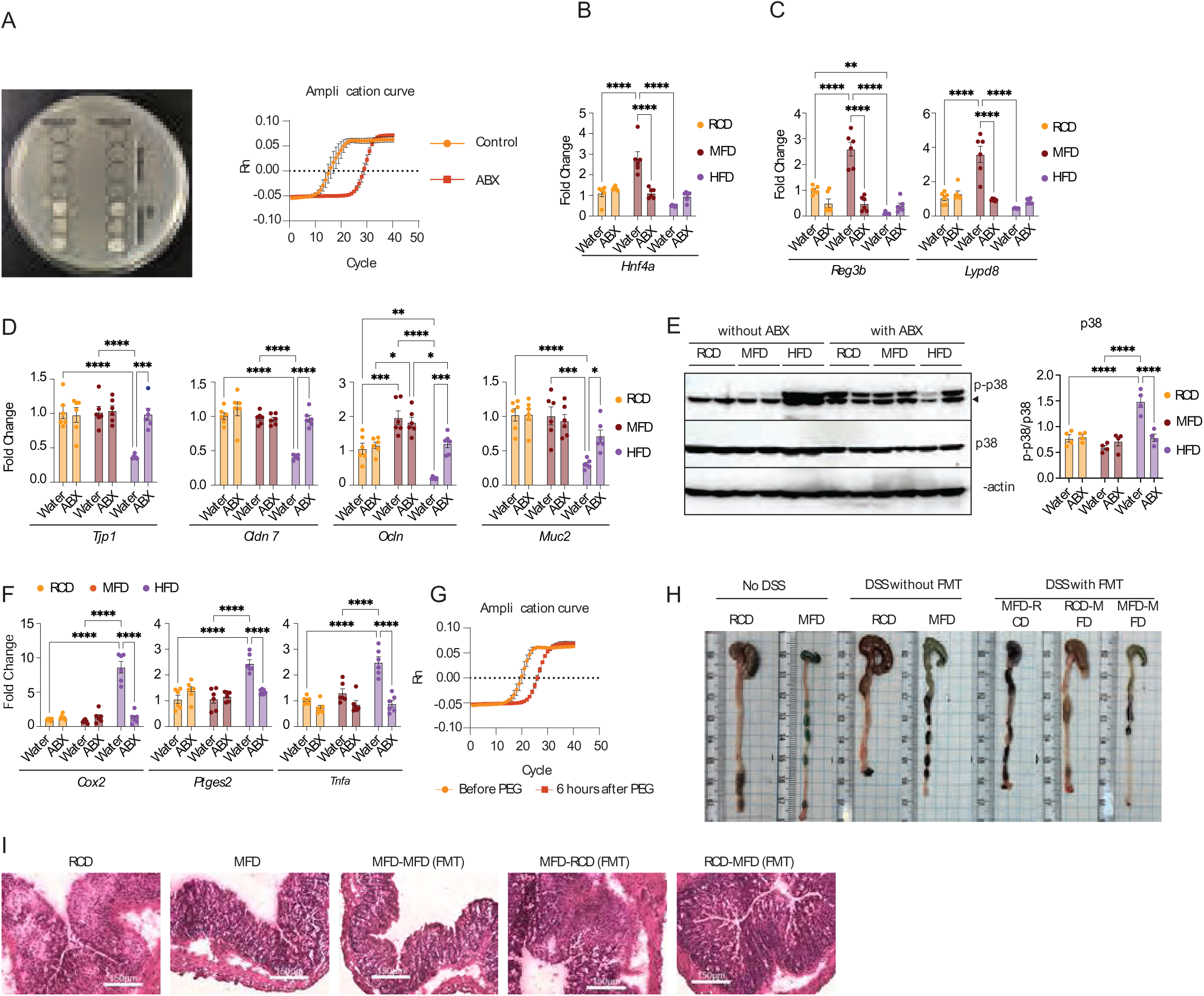
The gut microbiome directs the protective effects of milk based diets. **A:** LB agar plates showing fecal microbiota depletion in antibiotic-treated (ABX) mice; each black circle represents an individual mouse sample (left panel). qPCR amplification plot of the 16S rRNA gene from fecal DNA of ABX and control mice (n = 6) (right panel). **B,C,D&F:** RT-qPCR analysis of expression of AMPs associated genes (Hnf4α, Reg3b, Lypd8) (B,C), gut barrier genes (Tjp1, Cldn7, Ocln, and Muc2) (D), and inflammatory genes (Cox2, Ptges2, and Tnf) (F) in IECs of ABX-treated mice fed on RCD, MFD, and HFD (n = 6). **E:** Western blot analyses of IECs isolated from mice with or without antibiotic (ABX) treatment, fed RCD, MFD, and HFD, showing p-p38 and total p38 (n = 4). The panel on the right-hand side shows densitometric quantification. **G:** qPCR amplification plot of the 16S rRNA gene from fecal DNA before and 6 hours after PEG treatment (n = 6). **H:** Representative images of colon tissues from DSS or water-treated mice across various FMT groups. **I:** Representative images of H&E-stained colon tissue sections of various FMT groups with DSS. (Scale bars: 150 μm). Data are mean ± SEM of two independent experiments, with statistical analysis by two-way ANOVA with Tukey’s multiple comparison test (B, C, D, E, and F). *p < 0.05, **p < 0.01, ***p < 0.005, ****p < 0.001.

**Figure S7:**
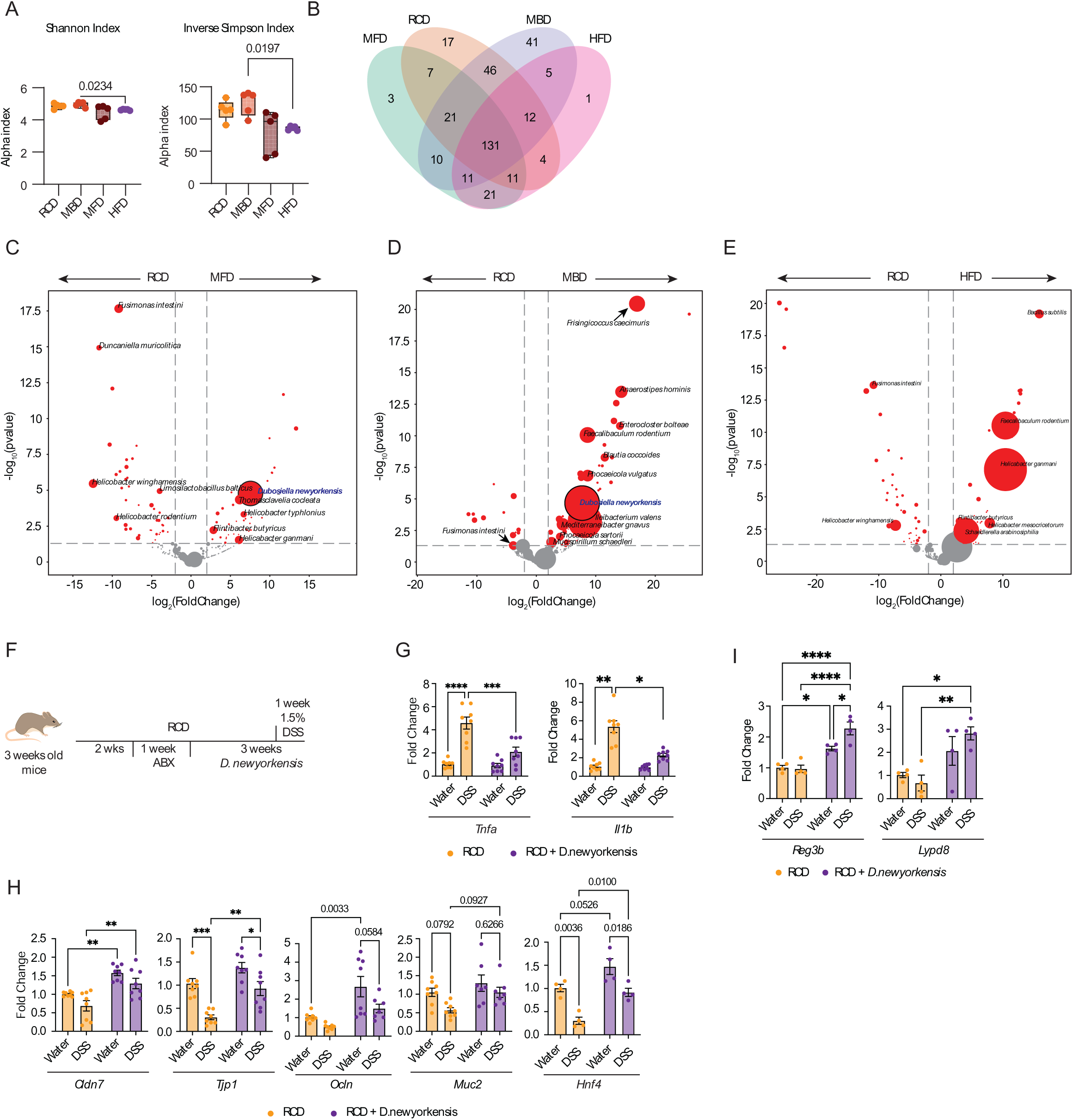
D. newyorkensis provides protective effect against DSS induced inflammation. **A:** Microbial alpha-diversity index, Shannon and inverse Simpson index of fecal microbiome isolated from mice fed on RCD, MBD, MFD, and HFD (n = 4). **B:** Venn diagram illustrating the common and unique species in the microbiomes of RCD, MBD, MFD, and HFD-fed mice. **C-E:** Volcano plots showing differentially enriched species across the conditions: MFD vs RCD (C), MBD vs RCD (D), and HFD and RCD (E). The size of the dots indicates the abundance of the species. **F:** Schematic illustration of the experimental workflow for D. newyorkensis treatment. **G-I:** RT-qPCR analysis of proinflammatory genes (*Tnfα,* and *Il1β*) (G), gut barrier genes (*Cldn7, Tjp1, Ocln, Muc2,* and *Hnf4α*) (H), and antimicrobial peptides (*Reg3b,* and *Lypd8*) (I) in colon tissues of D. newyorkensis supplemented mice with or without DSS treatment (n = 8). Data are mean ± SEM, with statistical analysis by Kruskal-wallis test with Dunn’s multiple comparisons test (A), DESeq2 (Wald test) Padj < 0.01 and absolute Log2 foldchange of > 1 (C, D, and E), and two-way ANOVA with Tukey’s multiple comparison test (G,H, and I). *p < 0.05, **p < 0.01, ***p < 0.005, ****p < 0.001.

**Figure S8:**
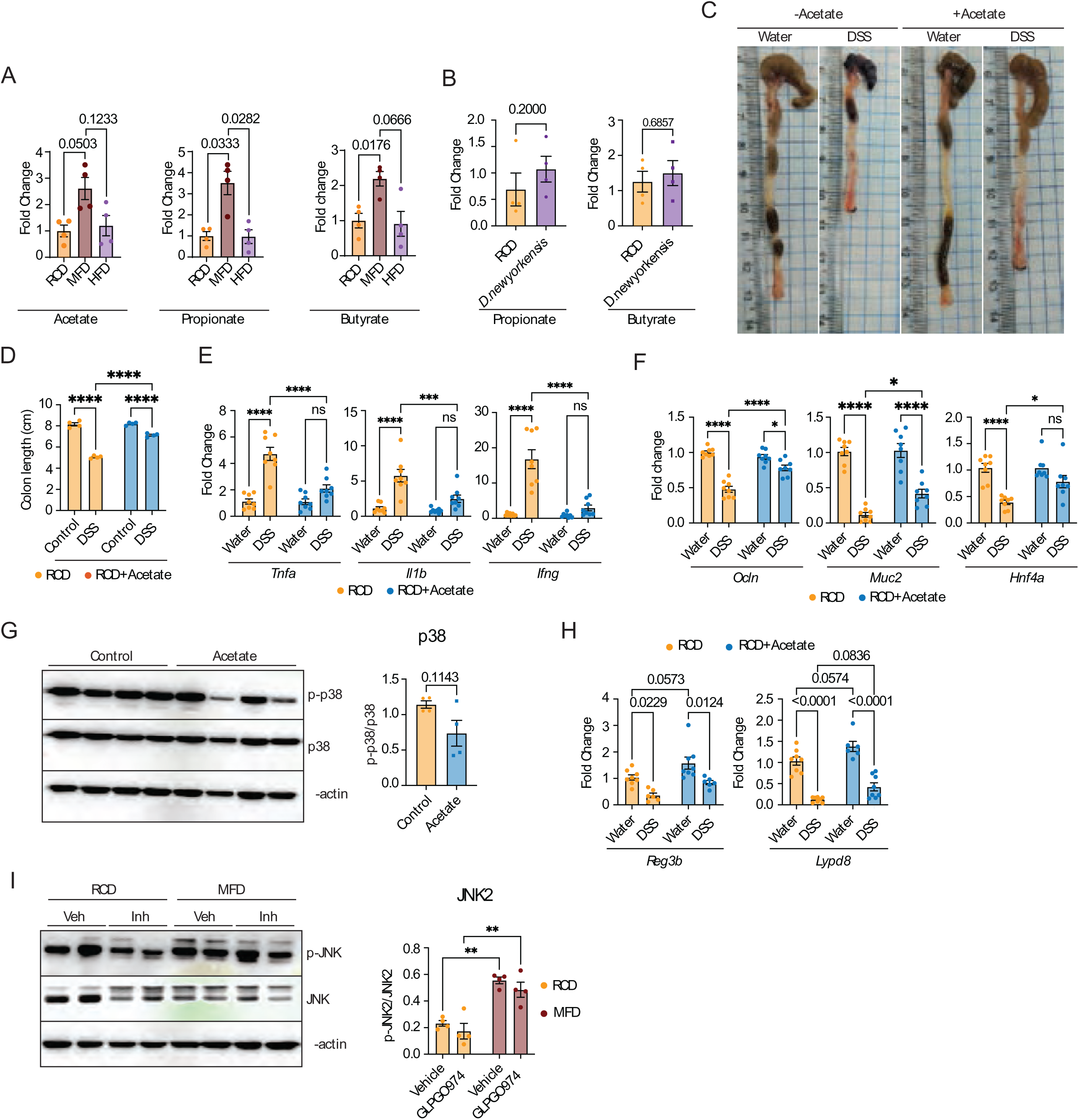
Acetate activates JNK2 and provides protection against DSS induced inflammation. **A:** Quantification of fecal matter acetate, propionate, and butyrate using GC-MS in different experimental groups (n = 4). **B:** Levels of propionate, and butyrate in the fecal matter of mice treated with D. newyorkensis and control mice (n = 4). **C&D:** Representative image of colon tissues (C) and total colon length in cm (D) of acetate and DSS-treated mice (n = 4). **E-F&H:** RT-qPCR analysis of proinflammatory cytokines (T*nfα, Il1β,* and *Ifnγ*) (E), gut barrier genes (*Ocln, Muc2,* and *Hnf4α)* (F), and AMPs (*Reg3b,* and *Lypd8*) (H) in colon tissues of mice treated with acetate with or without 1.5% DSS challenge (n = 8). **G:** Western blot analyses of IECs isolated from control and acetate-treated mice showing p-p38 and total-p38 levels (n = 4). Densitometric quantitation is given in the right panel. **I:** Western blot analyses of IECs isolated from GPR43 inhibitor (GLPG0974), and control mice showing p-JNK2 and total-JNK2 levels (n = 4). Densitometric quantitation is given in the right panel. Data are mean ± SEM, with statistical analysis by Brown-Forsythe and Welch ANOVA test (A), Mann Whitney test (B), two-way ANOVA with Tukey’s multiple comparison test (D, E, F, H, I, and J), and Unpaired T test with Welch’s correction (G). *p < 0.05, **p < 0.01, ***p < 0.005, ****p < 0.001.

**Figure S9:**
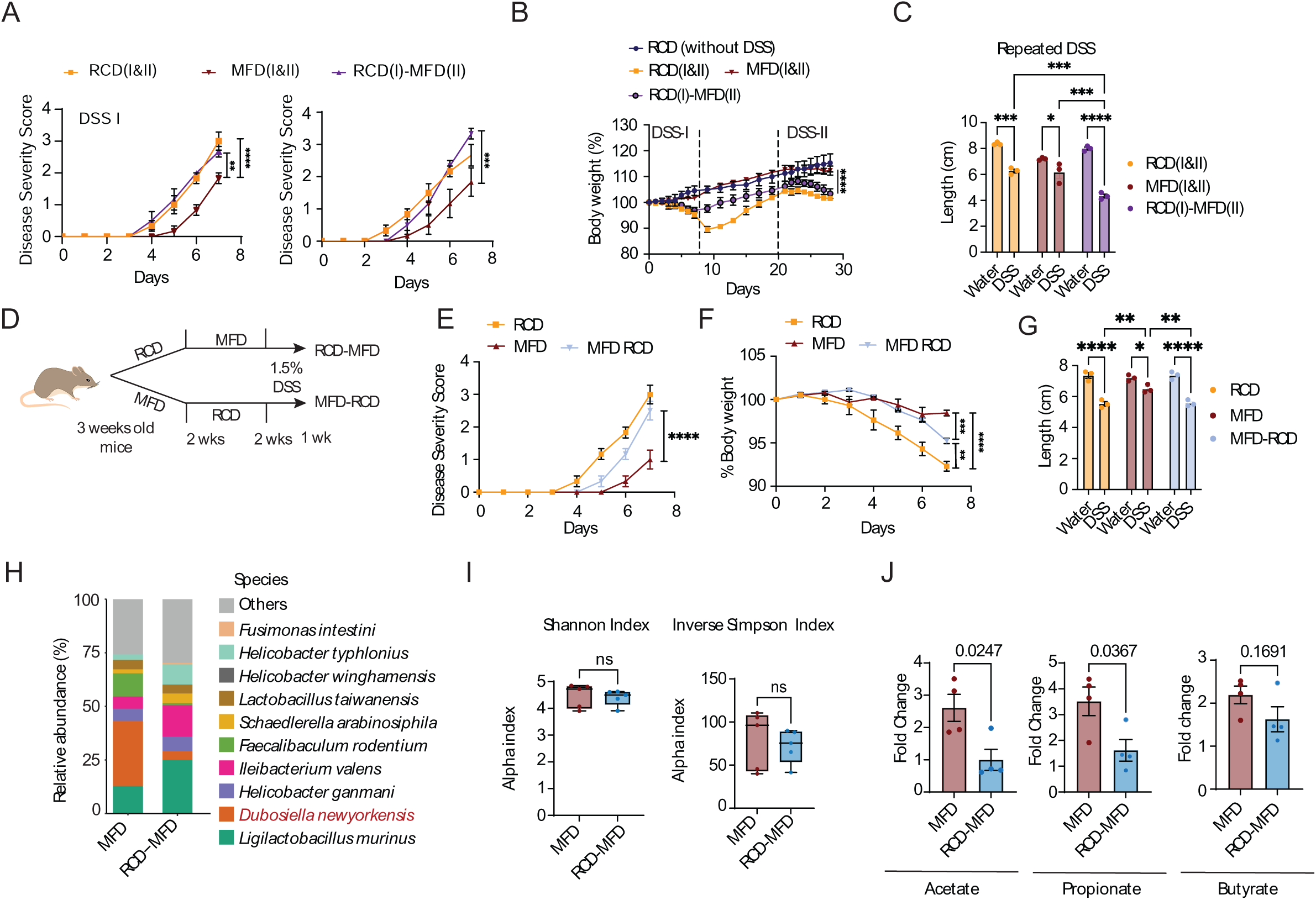
Dietary intervention during the weaning period is essential for establishing D. newyorkensis. **A:** Disease severity scores of RCD(I&II), MFD(I&II), and RCD(I)-MFD(II) groups of mice treated with 1.5% DSS, displaying scores from both rounds of the DSS insults (DSS-I & DSS-2) (n = 3). **B:** Daily body weight percentage (relative to Day 0) over 28 days in RCD(I&II), MFD(I&II), and RCD(I)-MFD(II) groups, with or without 1.5% DSS treatment (n = 3). Dashed vertical lines indicate periods of DSS administration. **C:** Colon length of RCD(I&II), MFD(I&II), and RCD(I)-MFD(II) groups of mice treated with two rounds of DSS or water (n = 3). **D:** Schematic illustration of the experimental workflow for data shown in Figures 5 F-P. **E-G:** Disease severity scores (E), body weight change (F), and total colon length in cm (G) of RCD, MFD, and MFD-RCD mouse groups, with or without DSS treatment (n = 3). **H:** Stacked bar plot showing the relative abundance of the top 10 microbial species in MFD and RCD-MFD groups, D. newyorkensis is highlighted. **I:** Microbial alpha-diversity index, Shannon and inverse Simpson index of fecal microbiome isolated from mice fed on RCD-MFD, and MFD (n = 5). **J:** Quantification of fecal acetate, propionate, and butyrate in MFD and RCD-MFD groups using GC-MS (n = 4). Data are mean ± SEM, with statistical analysis by two-way ANOVA with Šídák’s multiple comparison test (A, B, E, and F), two-way ANOVA with Tukey’s multiple comparison test (C, and G), Mann Whitney test (I), and Unpaired T test with Welch’s correction (J). *p < 0.05, **p < 0.01, ***p < 0.005, ****p < 0.001.

