## Supplementary Tables for "The diet-microbiome axis instructs gut barrier integrity by modulating colonocyte JNK-P38 duality"

**Table S1**

| | Chow | | MBD | | MFD | | HFD | |
|---|---|---|---|---|---|---|---|---|
| | Altromin International | | Nestle (Lactogen 2) | | Research Diets | | Research Diets | |
| | gm% | kcal% | gm% | kcal% | gm% | kcal% | gm% | kcal% |
| Protein | 23.7 | 30 | 14 | 12 | 26 | 20 | 26 | 20 |
| Carbohydrate | 38 | 50 | 59 | 50 | 26 | 20 | 26 | 20 |
| Fat | 7 | 20 | 20 | 38 | 35 | 60 | 35 | 60 |
| Energy Density | 333.9 kcal/100g | | 469 kcal/100g | | 523.73 kcal/100g | | 521.2 kcal/100g | |

**Table S1: Macronutrient composition of the diets used in this study**.

**Table S2**

| Sl. No | Module | Gene list |
|---|---|---|
| 1 | Inflammation associated genes | *Ifng, Ifngr1, Ifngr2, Il10,Il12a, Il12b, Il12rb1, Il12rb2, Il13, Il17a, Il17f, Il18, Il18r1, Il18rap, Il1a, Il1b, Il2, Il21, Il21r, Il22, Il23a, Il23r, Il2rg, Il4, Il4r, Il5, Il6, Jun, Nfkb1, Rela, Rora, Rorc, S100a8, S100a9, Stat1, Stat3, Stat4, Stat6, Tgfb1, Tgfb2, Tgfb3, Tnf* |
| 2 | Gut barrier repair gene set | *Aqp8, Sult1a1, Hsd17b2, Padi2, Slc26a2, Selenbp1, Fam162a, Acads, Tst* |

**Table S2: List of genes used to calculate module scores.**

| Table S3: Chemicals and Reagents | | |
|---|---|---|
| **REAGENT or RESOURCE** | **SOURCE** | **IDENTIFIER** |
| AMPICILIN SODIUM SALT | Sigma-Aldrich | A9518 |
| AMPure® XP beads | Beckman Coulter | A63881 |
| Anhydrous Sodium sulphate | Merck | 17521 |
| Brucella Agar base with Hemin | Himedia Laboratories | M1039 |
| BSTFA | Merck | B-023 |
| Cell strainer pore size 40um | Himedia Laboratories | TCP024 |
| Cell strainer pore size 70um | Himedia Laboratories | TCP025 |
| Chemiluminescent Substrate | Genaxy | XLS070,0250 |
| Collagenase IV | ThermoFisher | 17104-019 |
| Dextran sulfate sodium salt | MP Biomedicals | 160110 |
| Diethyl ether | Merck | 346136 |
| Dispase | Himedia Laboratories | TC303-100MG |
| DL-Dithiothreitol | Sigma-Aldrich | 43819 |
| EDTA, disodium salt | Bio Basic | EB0185-500 |
| FBS | Gibco | 10270-106 |
| FITC-Dextran 4 | Sigma-Aldrich | 46944 |
| Fluoroshield | Sigma-Aldrich | F6057 |
| GLPG0974 | Sigma-Aldrich | SML2443 |

| | | |
|---|---|---|
| HEPES | Sigma-Aldrich | 7365-45-9 |
| METRONIDAZOLE | Sigma-Aldrich | M3761 |
| Modified Tryptone Glucose Meat extract (MTGE) broth | Himedia Laboratories | M1116 |
| NEOMYCIN SULPHATE | Sigma-Aldrich | N6386 |
| JNK inhibitor II | Sigma-Aldrich | SP600125 |
| PBS | Gibco | 70011069 |
| PEG 6000 | Sysco | 49194 |
| Pierce BCA Protein Assay Kit | ThermoFisher | 23225 |
| PolyFreeze Tissue Freezing Medium | Sigma-Aldrich | SHH0026 |
| Potassium acetate | MP Biomedicals | 02194843-CF |
| Protease/Phosphatase Inhibitor | Cell signaling | 5872 |
| PVDF Transfer Membranes | ThermoFisher | 88520 |
| RIPA lysis buffer | Sigma-Aldrich | R0278 |
| RPMI 1640 medium | Gibco | 61870036 |
| SUCROSE | Sigma-Aldrich | S0389 |
| 2,4,6-trinitrobenzenesulfonic acid (TNBS) | Sigma-Aldrich | P2297 |
| Tween 20 | Sigma-Aldrich | 8.22184.0521 |
| VANCOMYCIN HYDROCHLORIDE | Sigma-Aldrich | V2002 |

Table S3: List of chemicals and reagents used in the study.

| Table S4: Antibodies | | | |
|---|---|---|---|
| **Western Antibodies** | | | |
| **REAGENT or RESOURCE** | **SOURCE** | **IDENTIFIER** | **DILUTION** |
| Cleaved caspase 3 | CST | 9664T | 1/1000 |
| Cyclin D1 | CST | 2978 | 1/1000 |
| p38 Antibody | CST | 8690 | 1/1000 |
| phospho-p38 Antibody | CST | 4511 | 1/1000 |
| MKK4 | CST | 9152 | 1/1000 |
| MKK7 | CST | 4172 | 1/1000 |
| Occludin (OC-3F10) | Thermo Fisher Scientific | 33-1500 | 1/1000 |
| SAP/JNK Antibody | CST | 9252 | 1/1000 |
| phospho-SAP/JNK Antibody | CST | 4668 | 1/1000 |
| Ki67 | CST | 9129 | 1/400 |
| Alexa Fluor™ 568 | Thermo Fisher Scientific | A11036 | 1/2000 |
| Beta-actin Antibody | CST | 4967 | 1/1000 |
| Peroxidase AffiniPure™ Goat Anti-Rabbit IgG (H+L) | Jackson ImmunoResearch | AB_2313567 | 1/10000 |

| | | | |
|---|---|---|---|
| AffiniPure Anti-Mouse IgG (H+L) Fab Fragment Goat Secondary Antibody | Jackson ImmunoResearch | AB_2338476 | 1/10000 |

**FACS Antibodies**

| REAGENT or RESOURCE | SOURCE | IDENTIFIER | CLONE | DILUTION |
|---|---|---|---|---|
| FITC anti-mouse CD326 (Ep-CAM) Antibody | BioLegend | 118207 | G8.8 | 1/200 |
| APC anti-mouse CD45 Recombinant Antibody | BioLegend | 157606 | QA17A26 | 1/200 |
| PE anti-mouse CD19 Antibody | BioLegend | 152408 | 1D3/CD19 | 1/200 |
| PE/Cyanine7 anti-mouse CD3 Antibody | BioLegend | 100220 | 17A2 | 1/200 |
| TruStain FcX™ PLUS (anti-mouse CD16/32) Antibody | BioLegend | 156603 | S17011E | 1/1000 |

Table S4: List of antibodies used in the study.

| Table S5: Commercial kits | | |
|---|---|---|
| **REAGENT or RESOURCE** | **SOURCE** | **IDENTIFIER** |
| NucleoSpin RNA, Mini kit for RNA purification | Macherey-Nagel | 740955.5 |
| Primescript cDNA synthesis kit | TAKARA | 6110A |
| SYBR® Premix Ex Taq | TAKARA | RR420A |
| Pierce BCA Protein Assay Kit | ThermoFisher | 23225 |
| Single Cell 3' v3.1 Gel Beads kit | 10x Genomics | 2000164 |
| 16S barcoding kit 1-24 | Oxford-Nanopore Technologies | SQK-16S024 |
| QIAamp® Fast DNA Stool Kit | Qiagen | 51604 |
| Qubit™ dsDNA HS and BR Assay Kits | ThermoFisher | Q32854 |

Table S5: List of commercially available kits used in the study.

| Table S6: Experimental models: Organisms/strains | | |
|---|---|---|
| **REAGENT or RESOURCE** | **SOURCE** | **IDENTIFIER** |
| *Duboisiella newyorkensis* | ATCC | TSD-64-0.5ML |
| Conventional C57BL/6 mice | The Jackson Lab | cat 000664 |

Table S6: Use of live organisms used in the study.

| Table S7: Diets | | |
|---|---|---|
| **REAGENT or RESOURCE** | **SOURCE** | **IDENTIFIER** |
| Lactogen 2 | Nestle | |
| Milk Fat Diet | Research Diets | D19112203 |
| High Fat Diet | Research Diets | D12492I |
| Regular Chow Diet | Altromin | 1324 |

Table S7: List of rodent diets used in the study.

| Table S8: Software and algorithms | | |
|---|---|---|
| **REAGENT or RESOURCE** | **SOURCE** | **IDENTIFIER** |
| AxioVs40 (V 4.8.2.0) | ZEISS | https://www.micro-shop.zeiss.com |
| CellRanger Ver: 7.1.0 | 10x Genomics | https://www.10xgenomics.com |
| EPI2ME | Oxford Nanopore | https://nanoporetech.com |
| FlowJo v10.9.0 | BD Bioscience | https://www.flowjo.com |
| Graphpad Prism v10 | GraphPad Prism Inc | https://www.graphpad.com |
| ImageJ (v. 2.0.0-rc-43/1.51k) | NIH | https://imagej.net |
| Image-Pro Plus 6.0 | Media Cybernetics | https://image-pro-plus.software.informer.com |
| MSD ChemStation E.02.01.1177 | Agilent | https://www.agilent.com |
| QuantStudio Real-Time PCR software v1.3 | Thermo Fisher | https://www.thermofisher.com |
| R v4.2.2 | R | https://www.r-project.org |

Table S8: List of software used in the study.
